# Fungal volatiles affect lateral root formation and increase plant tolerance to osmotic stress

**DOI:** 10.64898/2026.10.28.762544

**Authors:** Esperanza Miñambres, José Fermoso-Domínguez, Sandra Rodríguez-Sufuentes, Pedro Marco-Montori, Óscar Lorenzo, Mónica Calvo-Polanco

## Abstract

Symbiotic fungi communicate with plants via volatile organic compounds (VOCs), even before physical contact. While fungal VOCs are known to promote plant development under control conditions, their effects under abiotic stress remain poorly characterized. Here, we exposed *Arabidopsis thaliana* seedlings to VOCs from the ectomycorrhizal fungi *Laccaria bicolor* and *Hebeloma cylindrosporum*, and the endophyte *Serendipita indica*, under osmotic stress. Time-course transcriptional analysis revealed a rapid, transient induction of auxin transport genes within 3 days of VOC exposure, with minimal early cytokinin modulation. Using auxin-(*aux1-7, axr1-3*) and cytokinin-resistant (*cre1, arr1-4*) mutants, we show that this early response bypasses classic auxin/cytokinin antagonism, promoting lateral root proliferation by day 10 under both control and stress conditions. Transcriptomic profiling indicated that fungal VOCs modulate glucosinolate and leucine metabolism and regulate potassium (*HAK5*) and iron (*IRT1*) transporters. Genetic and phenotypic analyses of *hak5-2* and *akt1-2* mutants, together with nitric oxide signaling components, revealed that potassium transport and nitric oxide-dependent IRT1 regulation are essential for translating VOC perception into developmental responses. These findings underscore the multifaceted mechanisms by which fungal VOCs coordinate plant development and abiotic stress tolerance.

**Highlight:** Fungal volatile organic compounds reprogram auxin and cytokinin signaling, ion transporters, and defense metabolism to enhance root development and osmotic stress tolerance in Arabidopsis seedlings.

## Introduction

The development of lateral roots is critical for root system architecture in higher plants, enabling efficient nutrients and water uptake from the soil (Santos Teixeira & Tusscher, et al. 2019). Plant hormones and environmental signals coordinate lateral root formation (Rawat & Laxmi, 2024), where auxins and cytokinins (CKs) act antagonistically through a dynamic crosstalk to coordinate lateral root development (Nenadić & Vermeer, 2021). Auxin transport occurs both cell-to-cell, through specific influx (AUX1 and LAX1) and efflux (PIN) carriers, as well as via long-distance movement through the phloem, with additional contributions from ATP-binding cassette transporters (ABCB). Auxin perception relies on TIR1/AFB receptors and AUX/IAA repressors, triggering proteasome-mediated degradation to release auxin response factors (ARFs) that drive developmental transcriptional responses (Kazan, 2013; Quint & Gray, 2006). Conversely, cytokinin (CK) signaling operates via a two-component phosphorelay where AHK histidine kinase receptors (AHK2, AHK3, AHK4/CRE1) transfer signals through AHPs to activate response regulators, including type-A ARRs (negative regulators) and type-B ARRs (transcription factors) (El-Showk et al., 2013; Werner & Schmülling, 2009). Under environmental constraints such as drought or high salinity, osmotic stress profoundly alters root development by disrupting the fine-tuned equilibrium between auxin and cytokinin pathways (Mallick et al., 2022). Specifically, osmotic stress triggers spatial shifts in auxin redistribution, perception, and signaling (partially mediated by key influx carriers like AUX1) which are critical to fine-tune primary root elongation and control lateral root emergence as an adaptive response (Naser & Shani, 2016). Concurrently, modulated cytokinin signaling, involving receptors such as AHK3 and downstream ARR regulators, acts alongside auxin gradients to dynamically integrate stress perception with root developmental plasticity, ultimately dictating root system architecture optimization under adverse osmotic conditions (Mallick et al., 2022; Naser & Shani, 2016).

Beneficial soil fungi that form mutualistic associations with plants are known to favour water and nutrient uptake (Fresno et al., 2023; Calvo-Polanco et al., 2019) and to have a role in the tolerance of plants to different abiotic stresses (Valenzuela-Aragon et al., 2025; Zwiazek et al., 2019). Among them, the ectomycorrhizal fungi *Laccaria bicolor* and *Hebeloma cylindrosporum* are well-studied model species due to their symbioses with diverse tree hosts such as *Pinus* and *Populus spp.* in temperate and boreal ecosystems (Ruytinx et al., 2021; Marmeisse et al., 2004), while the root endophyte *Serendipita indica* associates with a broad range of terrestrial and aquatic plants (Tyagi et al., 2022). The communication between plants and fungi occurs previous to direct colonization through diverse biochemical signals, including small RNAs (Ledford et al., 2024), hormones (Pons et al., 2020), small peptides (Valmas et al., 2023), and volatile organic compounds (Plett et al., 2024). Volatile organic compounds (VOCs) are low-molecular-weight carbon compounds that readily evaporate and disperse through air, liquids, and porous soils. Fungal VOCs are chemically diverse, comprising alcohols, ketones, esters, alkenes, thiols, monoterpenes, and sesquiterpenes (El Jaddaoui et al., 2023; Inamdar et al., 2020). Composition and concentration of VOCs profiles depend on several factors, such as the substrate on which they grow, humidity, temperature, plant-fungus interaction time and fungal development stage (El Jaddaoui et al., 2023; Inamdar et al., 2020). Recent studies have shown that fungal VOCs stimulate plant growth (Guo et al., 2021; Venneman et al., 2020), promoting lateral root and root development (Chiu et al., 2022; Ditengou et al., 2015), biomass accumulation and photosynthetic capacity (Venneman et al., 2020), and that this beneficial effect is independent of the potential pathogenic or symbiotic effect of the fungi in direct contact with the plants (Miñambres et al., 2026), at least in the first stages of plant development.

Fungal VOCs from specific species, including *L. bicolor* (Ditengou et al., 2015) and *S. indica* (Venneman et al., 2020), are known to affect auxin and/or CKs signaling, VOCs from *Laccaria bicolor* significantly enhance lateral root formation in *Arabidopsis thaliana* and *Populus tremula × alba* under controlled conditions by modulating the auxin signaling pathway, specifically through polar auxin transport mediated by PIN transporters (Ditengou et al., 2015). Sesquiterpenes, absent in *Cenococcum geophilum*, were identified as the key VOCs, with (–)-thujopsene inducing the greatest lateral root increase, suggesting its pivotal role in root development. Venneman et al. (2020), showed that the VOCs emitted by *Serendipita* spp. combined with different levels of CO_2_, stimulate *Arabidopsis* growth enhancing leaf expansion, root system development, and photosynthetic efficiency through both auxin and CKs signaling pathways. However, the predominant VOC from *Serendipit*a spp., methyl benzoate, could not replicate these effects. The importance of auxin and CKs pathways in microbial VOC-mediated plant growth is also supported by several studies on rhizobacteria (Pérez-Flores et al., 2017) and phytopathogens (Sánchez-López et al., 2016). Thus, understanding fungal VOCs signaling mechanisms is crucial for developing biotechnological strategies to improve crop and forest plants resilience, particularly in the context of climate change.

Our aim is to investigate the regulation of root development and the molecular effects of fungal VOCs on *Arabidopsis thaliana* seedlings under controlled and osmotic stress conditions. The plants will be exposed to the VOCs emitted by the non-host ectomycorrhizal fungi *Laccaria bicolor* and *Hebeloma cylindrosporum*, and the VOCs from the host-compatible root endophyte *Serendipita indica*.

**Table 1.** Selected compounds from the fungal volatile profile analyses.

| Compound | Type | Effect | References | Nº CAS | Commercial |
| --- | --- | --- | --- | --- | --- |
| 2-methyl-propanal | Aldehyde | Inhibition of cell development | Gupta et al., 2022 | 78-84-2<br>SIGMA | SIGMA |
| 3-octanone | Ketone | Through AUX/CK, it affects plant growth and lateral root development in *Streptomyces* | Dotson et al., 2020 | 106-68-3<br>SIGMA | SIGMA |
| Octanal | Aldehyde | Growth inhibitor of certain plants and antifungal agent | Hpoo et al., 2020; Li et al., 2021 | 124-13-0<br>SIGMA | SIGMA |
| Farnesylacetone | Ketone terpene | Effect on plant growth | Câmara et al., 2024; Groenhagen et al., | 762-29-8<br>SIGMA | SIGMA |
|  |  |  | 2013; Lee et al.,<br>2016 |  |  |

## Material and methods

### Biological material and growth conditions

*Arabidopsis thaliana* (L.) Heynh ecotype Columbia (Col-0) was the genetic background used in this work. All mutants used in this study are shown in Table 2. *Laccaria bicolor (Maire) P. D. Orton* strain S238O, was obtained from Dr. Francis Martin, INRAE Nancy, France, S238N (Bertaux et al., 2003; Di Battista et al., 1996). *Hebeloma cylindrosporum (Romagnesi)* obtained from Dr. Sabine Zimmerman, INRAE Montpellier. Endophytic fungi *Serendipita indica* Verma, Varma, Kost, Rexer & Franken (Varma et al., 1999; Verma et al., 1998) was obtained from Dr. Asunción Morte, University of Murcia. All the fungi were replicated every 30 days in Petri dishes with Pachlewski solid medium (Pachlewski & Pachlewska, 1974).

**Table 2.**
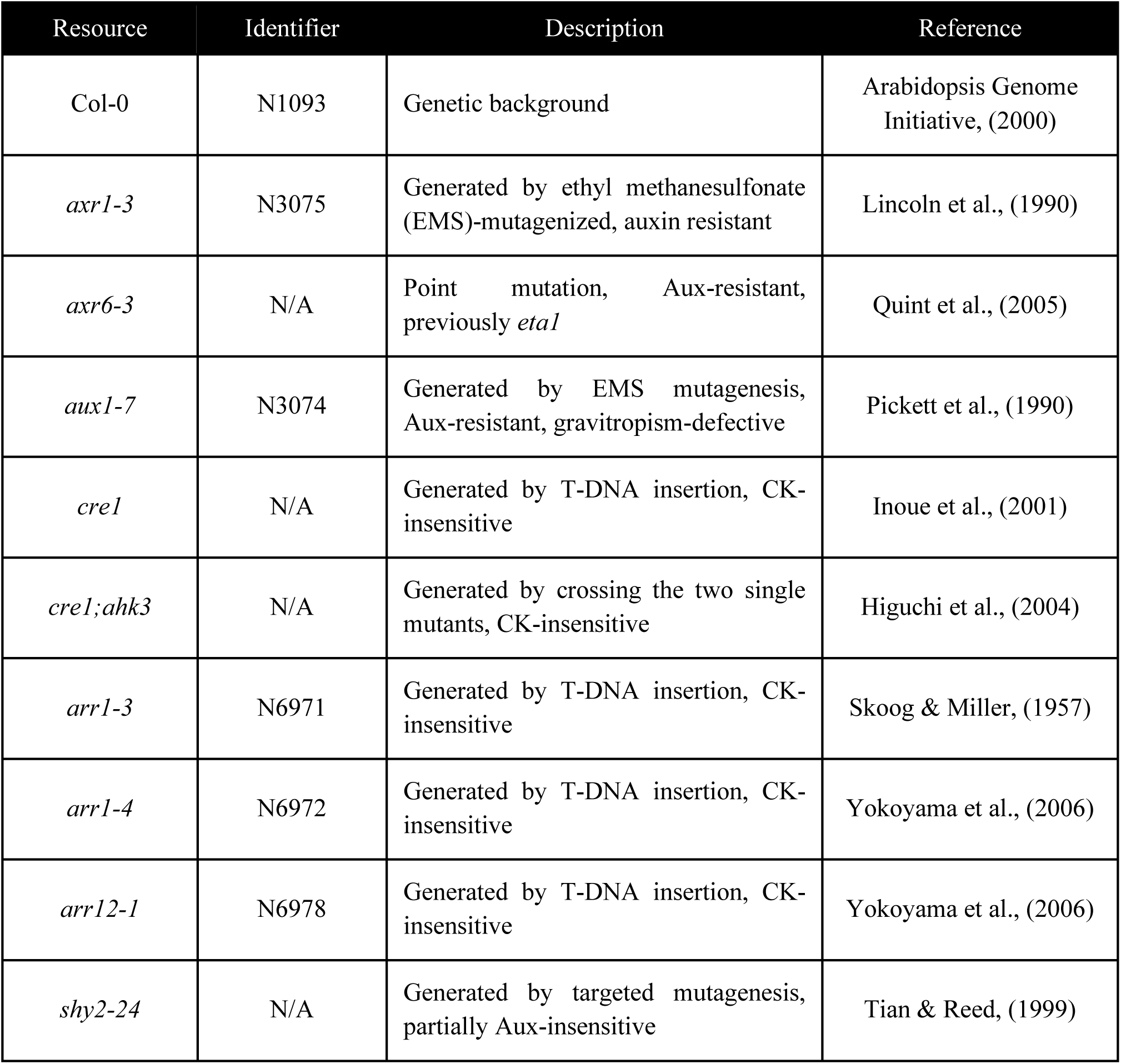
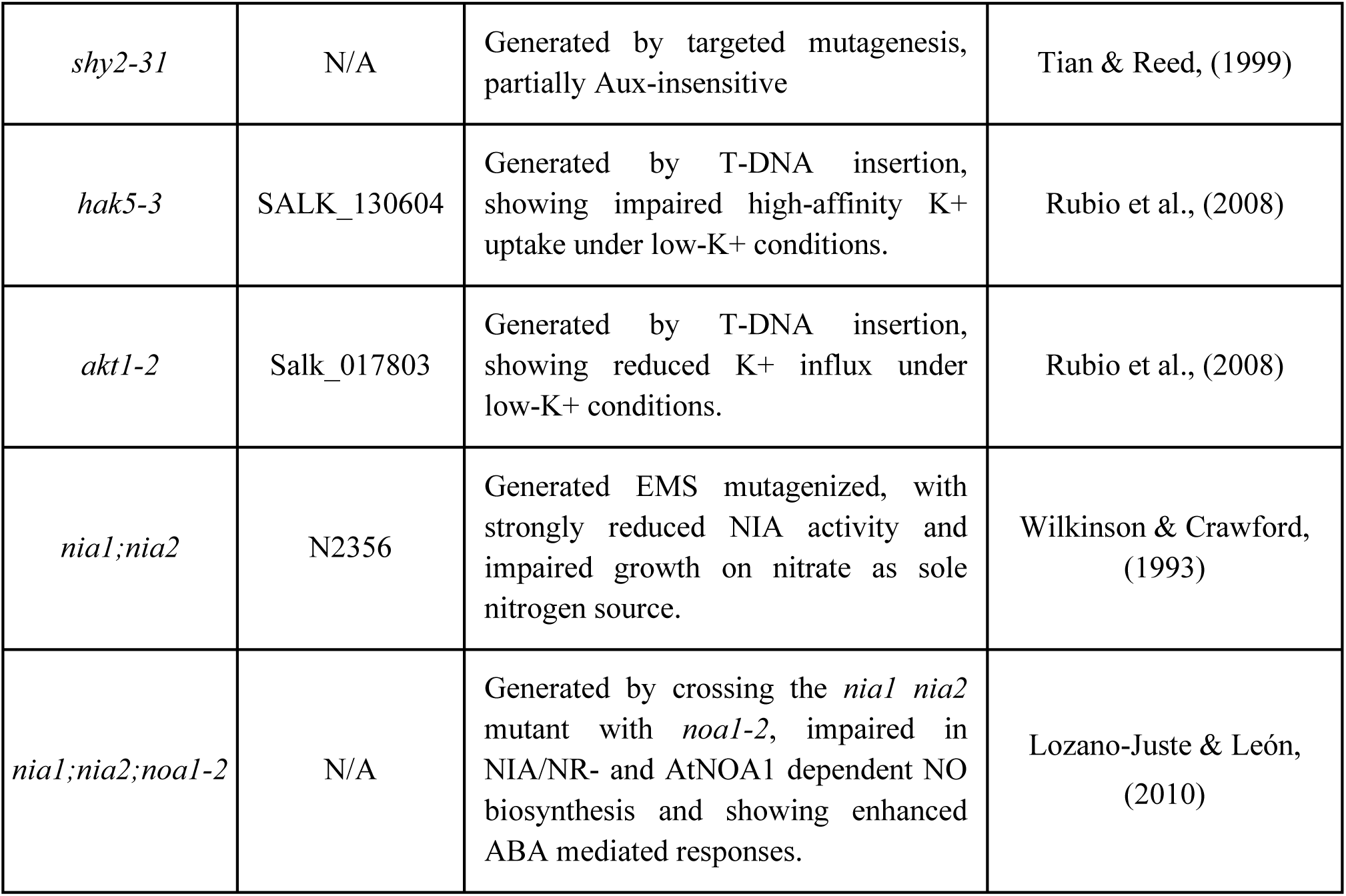
Arabidopsis lines used in this study including a brief description and the corresponding reference for each.

For the different experiments, Arabidopsis seeds were surface sterilized in 75 % (v/v) sodium hypochlorite and 0.01 % (v/v) Triton X-100 for 5 min and washed three times in sterile water before sowing. Seeds were stratified for 24 h at 4 °C. All plant-fungi co-cultives were grown in a Fitoclima 600 Aralab climate chamber under controlled conditions: 50-60% humidity, 16:8 h (day:night), 22:18 °C and light intensity of 150 μE m^−2^ s^−1^.

### Screening of *Arabidopsis thaliana* mutants of different hormonal pathways exposed to fungal VOCs under control conditions

For the screening experiments, four round Petri dishes (55 mm diameter, Corning) were placed into a 140 mm round Petri dish (Deltalab). Three of the 55 mm plates were filled with MS medium (Murashige, 1962) and one with Pachlewski P20 medium (P20, Pachlewski & Pachlewska, 1974). The P20 plate was used to grow the fungi (*L. bicolor*, and *S. indica*), and the MS plates were used to sow the Arabidopsis seeds. Sixty seeds of Col-0 or the Arabidopsis mutants were sown on each 55 mm plate and three replicates were performed for each genotype and condition (Supplementary Figure 1A).

Plates were scanned (EPSON V850 Pro) 6 and 10 days after sowing. The area of the cotyledons was measured at day 6 using *ImageJ* (Schneider et al., 2012), and the total fresh weight and chlorophyll content were determined at day 10.

For chlorophyll content, three independent biological replicates per genotype and condition were performed. For each biological replicate, five seedlings were pooled from a single plate and extracted with 100% methanol for 20 min. Absorbance was measured at 652 nm and 665 nm in a Varioskan LUX spectrophotometer (Thermo Scientific). Chlorophyll *a*, *b*, and total chlorophyll content were calculated as in Lichtenthaler (1987), normalized to total extract volume, and expressed as µg pigment mg⁻¹ fresh weight.

### Plant-fungus co-culture for phenotypic analysis of auxins, CKs, NO biosynthesis and ion transporter mutants under control and osmotic conditions

For the plant-fungus co-culture, Arabidopsis seeds were sawn into square plates (120×120mm, Fisher Scientific) filled with MS medium. Three round 35 mm plates (FALCON, A Corning Brand) were set into the square plates filled with P20 medium and were used to grow either *L. bicolor, S. indica* or *H. cylindrosporum.* Each square plate contained seeds of the wild-type Col-0 and one mutant genotype. For each genotype and condition, five independent biological replicates were performed, each consisting of one square plate with 20 seeds per genotype (Supplementary Figure 1B). The fungi were grown for 6 days before being inserted into the co-culture system. The square plates were covered with a black cover for the root and fungi to be in darkness.

For the osmotic stress condition, the plates were prepared in the same way as explained above but with MS medium supplemented with 100 mM sorbitol. Osmotic stress generates a lower water potential between the medium and the plant, making it difficult to take up water and thus resembling the water stress that plants might face if they were in the environment (Martínez-Santos et al., 2021).

When the seedlings were 10-day-old, the plates were scanned using an EPSON V850 Pro scanner, and their fresh weight and length of the primary and secondary roots were determined using *Image J* (Schneider et al., 2012).

### Analysis of the fungal VOCs profile

Fungal volatile compounds were analyzed from the headspace of cultures grown for 10 days on Pachlewski P20 agar plates (Pachlewski & Pachlewska, 1974) using PTR-TOF-MS (Proton Transfer Reaction-Time of Flight-Mass Spectrometry). an ultrasensitive analytical technique for real-time detection of VOCs (Ghaffari et al., 2025), at CARTIF Research Center (Parque Tecnológico de Boecillo, Valladolid, Spain). Air samples from 10-day-old fungal cultures were continuously monitored by introducing the sampling capillary directly into closed 35-mm Petri dishes containing fungal cultures for 3 minutes per sample. Raw data were processed using Ionicon Data Analyzer (IDA) software and further analyzed in Microsoft Excel.

To determine the metabolites fixed in the medium after 6 days of fungal growth at the square co-culture system, 1 g of agar from the MS medium was sampled and analyzed by the Elemental, Chromatography and Mass Analysis Facility (NUCLEUS, University of Salamanca). Agar from a plate not exposed to VOCs was compared with agar from plates exposed to VOCs to eliminate compounds normally present in the medium.

### Application of commercial VOCs to *Arabidopsis thaliana* plants

Based on the fungal volatiles identified in the fungal VOC profile analyses, the following compounds were selected: farnesylacetone, 2-methyl-1-propanal, octanone and octanal. These were purchased in commercial formulation (Thermo Scientific) and a series of dose-response tests were carried out to identify concentrations that balanced phenotypic effects with minimal phytotoxicity. The concentrations of 62.5 mM, 125.0 mM, and 250.0 mM were selected for farnesylacetone and 2-methyl-1-propanal; 7.8 mM, 15.6 mM, and 31.3 mM for octanone; and 31.25 mM, 62.5 mM, and 125.0 mM for octanal.

Wild type (Col-0) plants were grown in the 120×120 mm square co-culture system, but in this case pre-sterilized filter paper was placed on the 35 mm plates (FALCON, A Corning Brand) and moistened with 200 µL of the compound or sterile distilled water (control). Arabidopsis seeds were sown after applying the corresponding compound to the filter paper. Twenty Arabidopsis seeds were sown per plate, and five independent biological replicates were performed (Supplementary Figure 1C).

Root length, number of lateral roots and fresh weight were measured after 10 days of exposure to the commercial volatiles. Root lengths were measured with *ImageJ* (Schneider et al., 2012).

### Root RNAseq profile

Total RNA was extracted from roots of 10-day-old seedlings. Briefly, frozen tissue was powdered with mortar and conserved at −80°C until use. RNA was extracted with RNeasy Plant Mini Kit (QIAGEN) and was precipitated in water RNAse free. RNA concentration was measured in NanoDrop 1000 (Thermo Scientific).

RNA integrity was evaluated using an Agilent 2100 Bioanalyzer (Agilent Technologies) with the 2100 Expert Eukaryote Total RNA Nano chip, prior to library construction. For each condition, three independent biological replicates were sequenced, each corresponding to RNA extracted from all roots harvested from a single plate. cDNA libraries were prepared using the Illumina Stranded mRNA Prep, Ligation kit (Illumina), following the manufacturer’s instructions. Libraries were quantified with the Qubit 4 Fluorometer (Thermo Fisher Scientific) using the dsDNA High Sensitivity Assay. Fragment size distribution and library quality were further assessed using the Agilent 2100 Bioanalyzer with the dsDNA High Sensitivity kit (Agilent Technologies). Library preparation and transcriptome sequencing were carried out using the NextSeq 1000/2000 system and P3 flow cell chemistry (200 cycles), generating 100 bp paired-end reads with a minimum of 25 million reads per sample.

The Illumina RNASeq raw reads were preprocessed using fastp (v0.20.0) (Chen et al., 2018) and the quality of the reads was estimated by FastQC v0.11.9 (http://bioinformatics.bbsrc.ac.uk/projects/fastqc). Clean reads were aligned against TAIR 10.1 GCF_000001735.4 reference genome using hisat2 v4.8.2 (Kim et al., 2015), and Samtools v0.1.19 was used to translate SAM file to BAM file and then to sort BAM files using default parameters (Li et al., 2009). FeatureCounts v2.0.1 in the subread package was used to generate read counts for each sample (Liao et al., 2014). Normalized gene expression levels, calculated by variance stabilizing transformation and differential expressed genes (DEGs) using q-value ≤ 0.05 and log_2_ (fold change) of < |1.0| as threshold were obtained using DESeq2 (Love et al., 2014). Volcano plots were generated by ggplot2 pac< | of R (Wickham, 2016). Venn diagrams were generated using *ggVennDiagram* package of R (Gao, et al. 2021). Heatmaps were generated using *pheatmap* package of R (Kolde, 2018). Gene ontology (GO) enrichment was performed on the differentially expressed genes, applying a p-value < 0.05 and a log2FoldChange > |1.0| and were generated by clusterProfiler of R (Yu, 2012), AnnotationDbi (Pagès, 2024) and org.At.tair.db (Carlson, 2024) and annotated genes of TAIR10 (Berardini, 2015). Bubble plots were generated using the ggplot2 package (Wickham, 2016).

### Time-course analysis auxin and cytokinin genes

Total RNA was extracted from whole seeds at 24 and 48 hours post-sowing (including emerging radicles when present), and from isolated roots of 3, 5, 7, and 10-day-old seedlings (calculated from sowing). Briefly, frozen tissue was ground to a fine powder using a mortar and pestle and stored at –80 °C until use. The homogenate was prepared in extraction buffer (0.2 M Tris-HCl pH 8.0, 25 mM EDTA, 0.4 M LiCl, 1% SDS, prepared in DEPC-treated water) with an equal volume of chloroform, vortexed, and centrifuged. The supernatant was sequentially extracted with phenol/chloroform (1:1, v/v) and precipitated with LiCl at –80 °C overnight. The RNA pellet was recovered by centrifugation, washed sequentially with 3 M sodium acetate (pH 5.2) and 100% ethanol, followed by a final wash with 70% ethanol. Pellets were dried using a SpeedVac, resuspended in DEPC-treated water, and stored at –80 °C. RNA concentration and purity were measured using a NanoDrop 1000 spectrophotometer (Thermo Scientific).

### Quantitative reverse transcriptase–PCR (RT-qPCR) analysis

To analyze the expression dynamics of auxin- and cytokinin-related genes across a time course and to validate candidate genes identified by RNA-seq analysis, reverse transcription was performed using the NZY First-Strand cDNA Synthesis Kit (NZYtech). Then, RT-qPCR was conducted on a StepOnePlus Real-Time PCR system (Thermo Fisher Scientific) using SYBR FAST Rox High qPCR master mix (KAPA Biosystems). The thermal cycling program consisted of 50°C for 2 min, 95°C for 10 min, followed by 40 cycles of 95°C for 15 s and 60°C for 1 min. Ct values were determined using StepOne Software version 2.3, and relative expression levels were calculated, normalized to the reference genes *Elongation Factor 1* (*EF1*) and *Ubiquitin 5* (*UBQ5*). Primers used are listed in Supplementary Table 1. Three independent biological replicates were performed, each analyzed in three technical replicates.

### Statistical analysis

Data on cotyledon areas, chlorophyll content, primary root length, number of lateral roots, and fresh weight were analyzed using one way ANOVA together with Tukey post-hoc test using R 4.4.2 (R Core Team, 2024). The normality and homoscedasticity of the data were tested for the assays. All graphics were generated by the ggplot2 package of R 4.4.2 (R Core Team, 2024; Wickham, 2016).

## Results

### Volatile organic compound profiles of Laccaria bicolor, Hebeloma cylindrosporum and Serendipita indica

VOCs emitted by fungi are highly diverse and their profile is influenced by numerous external factors (El Jaddaoui et al., 2023; Hung et al., 2015). VOCs emitted by *L. bicolor*, *H. cylindrosporum*, and *S. indica* were in total 172, 126 of them emitted by all three fungi (Figure 1A), with aldehydes, alcohols, esters, ketones, and aromatic compounds as the most represented ones. Moreover, heatmaps of relative VOC emission intensities (Figure 1B, Supplementary Figure 2A) revealed highly similar emission patterns across the three fungi species, with all fungi exhibiting parallel trends in compound abundance. On the other hand, the analyses of the VOCs present in the Murashige Skoog (MS) medium (Murashige, 1962) in the co-cultivation system detected a total of 45 compounds, where carboxylic acids, amides, and lactones were identified (Figure 1C).

**Figure 1.**
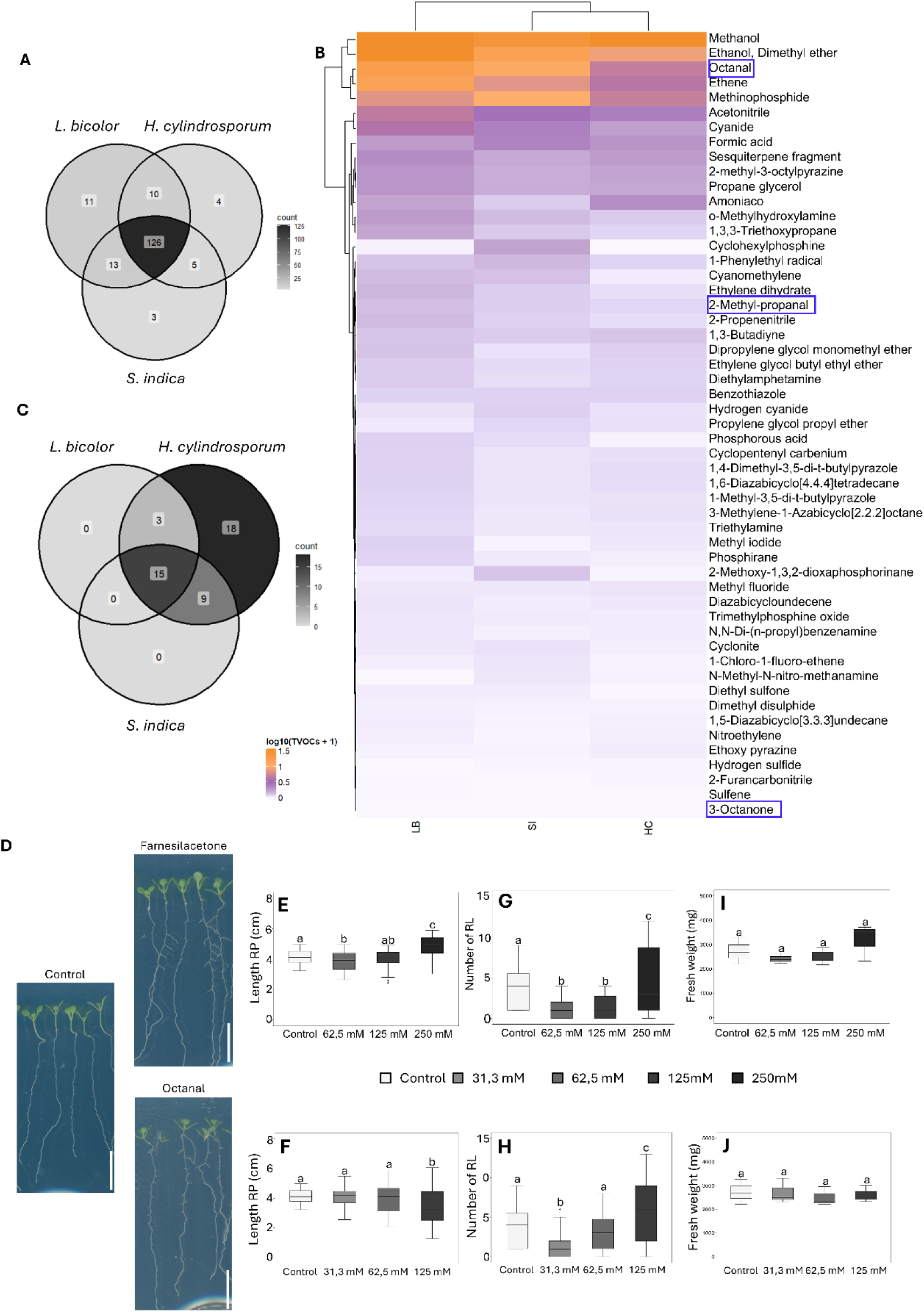
VOCs profile of the three fungi under study analyzed by HS-GC-MS and analysis of Col-0 seedling development at 10 days in the presence of farnesylacetone and octanal. A. Venn diagram shows that 126 of the compounds analyzed are common VOCs emitted by all three fungi. B. Heatmap of VOCs emitted under the three fungi. Color intensity indicates the relative abundance of each compound in each condition, with rows representing individual volatiles and columns representing treatments. C. Venn diagram showing common VOCs emitted by all three fungi. D. Phenotype of Col-0 seedlings, scale bar=1 cm. E-F. Primary root length (PRL) (cm). G-H. Number of lateral roots (LR). I-J. Total fresh weight of seedlings (mg). Data are shown as boxplots of five biological replicates (n=5 plates) per genotype and condition; each plate contained 20 seeds per genotype. Different letters indicate statistical differences (p<0.05) among treatment means using one-way ANOVA and Tukey’s test.

Four compounds, octanal, 2-methyl propanal, 3-octenone, and farnesylacetone, were selected for phenotypic testing (Table 1, Figure 1D-J Supplementary Figure 2B-G) as they were previously shown to affect plant development. Initial screening allowed the identification of farnesylacetone and octanal as the only compounds producing significant developmental changes (Figure 1D-J Supplementary Figure 2B-G). In particular, farnesylacetone at 250 mM and octanal at 125 mM increased primary root length (Figure 1E-F) and promoted lateral root formation (Figure 1G-H). These concentrations were therefore selected for further analysis. No significant changes in fresh weight were observed with any of the tested compounds (Figure 1I-J, Supplementary Figure 2F-G). The remaining compounds showed no significant effects on primary root length (Supplementary Figures 2B-C) or lateral root number (Supplementary Figures 2D-E). Having established that individual VOCs elicit only partial developmental effects, we next examined the impact of complete fungal volatile profile across hormonal mutant backgrounds.

### Effects of fungal VOCs in Arabidopsis fresh weights, cotyledons development and leaf chlorophyll contents under control conditions

A screening was conducted using wild type (Col-0) Arabidopsis seedlings and 17 mutants defective in cytokinin, auxin (Figure 2), ethylene, jasmonic acid, gibberellins, abscisic acid signaling/perception (Supplementary Figures 3–4). Seedlings were exposed to VOCs from *L. bicolor* and *S. indica*.

**Figure 2.**
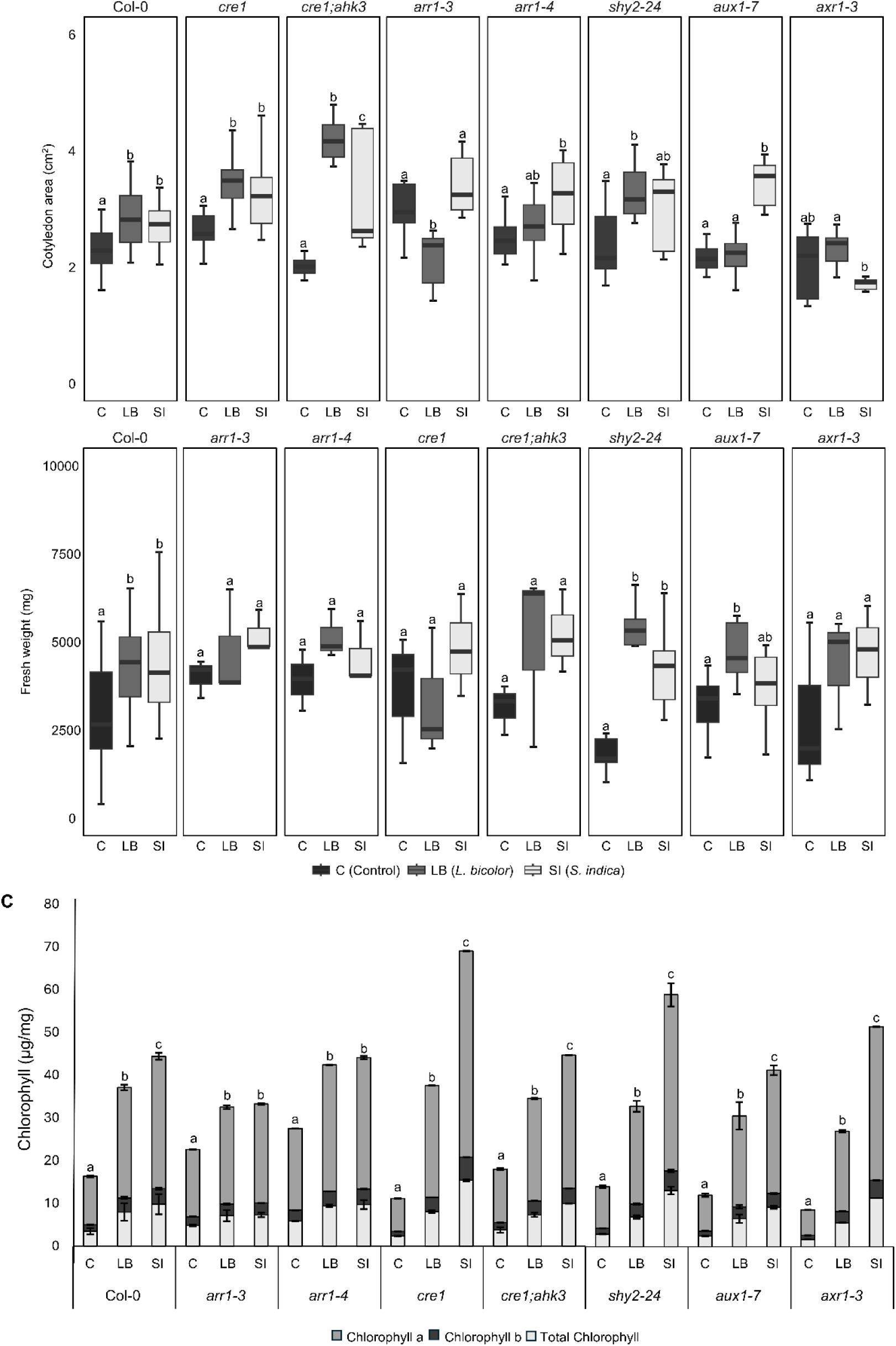
Analysis of seedling development of CK pathway mutants (*arr1-3*, *arr1-4*, *cre1*, *cre1;ahk3*) and auxin mutants (*shy2-24, aux1-7, axr1-3*) in the presence and absence of VOCs emitted by *L. bicolor* and *S. indica*. A. Cotyledon area of 6-day-old seedlings (cm^2^). B. Fresh weight of 10-day-old seedlings (mg). C. Total chlorophyll, chlorophyll a, and chlorophyll b (µg/mL) at 10 days-old seedlings. Three biological replicates (n=3 plates) per genotype and condition; each plate contained 60 seedlings. For chlorophyll measurements, five seedlings were pooled per biological replicate. Different letters indicate significant differences among treatment means according to one-way ANOVA followed by Tukey’s test (P < 0.05).

The area of the cotyledons in Col-0 seedlings increased in the presence of all fungal VOCs compared to the not exposed ones (Figures 2A). The CKs mutants *cre1* and *cre1;ahk3* showed an increase in cotyledon area only in the presence of *L. bicolor* VOCs, while they were reduced in the *arr1-3* CK mutant (Figure 2A). In the auxin mutants, *shy2-24* showed an increase in cotyledon area when subjected to the *L. bicolor* VOCs, while in the mutant *aux1-7* increased in the presence of *S. indica* VOCs (Figure 2A). This genotype-specific response suggests that auxin signaling (via SHY2) and transport (via AUX1) may be triggered by distinct volatile compounds present in the unique profile of each fungus, despite sharing a core set of VOCs (Figures 1A and 1C).

As for the seedlings fresh weight, the VOCs emitted by both *L. bicolor* and *S. indica* induced an increase in Col-0 and *shy2-24* seedlings (Figures 2B), while in the *aux1-7* mutant there was only significant increase in the presence of VOCs emitted by *L. bicolor* (Figure 2B). Total chlorophyll content increased in Col-0 and in all CKs and auxin mutants analyzed, both in the presence of VOCs emitted by *L. bicolor* and by *S. indica*, compared to the absence of fungal VOCs (Figures 2C).

These results reinforce the idea that auxin and cytokinin pathways are primary mediators of fungal VOC effects on Arabidopsis development under non-stressed conditions as shown in previous studies (Venneman et al., 2020; Ditengou et al., 2015).

### Fungal VOCs transiently modulate auxin and cytokinin pathway gene expression under control conditions

To decipher whether these hormonal pathways are actively regulated upon initial volatile perception or during later growth stages, we characterized the temporal transcriptional dynamics of key auxin- and cytokinin-related genes under control conditions. The expression of key auxin transport (*AUX1* and *EIR1*) and signaling (*SHY2/IAA3*) genes, as well as cytokinin (CK) biosynthesis (*IPT3*) and transport (*PUP14*) genes, was monitored by RT-qPCR at 24 h and 48 h (whole germinating seeds) and at 3, 5, 7, and 10 days post-sowing (dps; isolated roots) under continuous VOC exposure (Figure 3).

**Figure 3.**
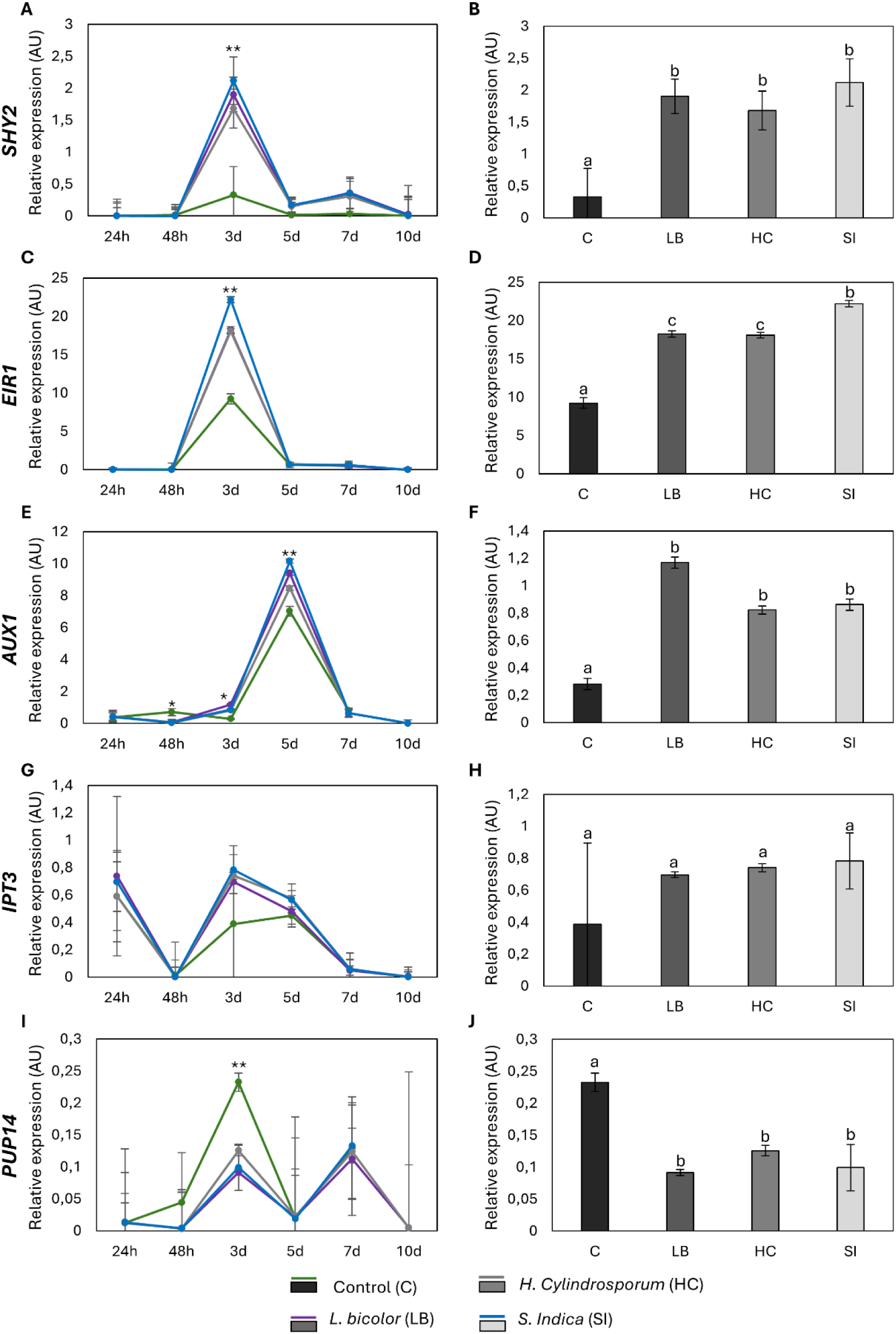
Analysis of the expression of key auxin transport (*AUX1* and *EIR2*) and signaling genes (*SHY2/IAA3*) and cytokinin biosynthesis (*IPT3*) and signaling genes (*PUP14*) by qRT-PCR at 3, 5, 7, and 10 days after exposure with fungal VOCs. A-B. Relative expression of *SHY2* over time and at 3 days. C-D. Relative expression of *EIR2* over time and at 3 days. E-F. Relative expression of *AUX1* over time and at 3 days. G-H. Relative expression of *IPT3* over time and at 3 days. I-J. Relative expression of *PUP14* over time and at 3 days. Data are mean ± SE of three biological replicates (n=3 independent plates), each analyzed in three technical replicates. Asterisks indicate statistically significant differences among treatment means (* p<0.05, ** p < 0.01) using one-way ANOVA and Tukey’s test. And different letters indicate significant differences among treatment means according to one-way ANOVA followed by Tukey’s test (P < 0.05).

Across the entire time course, the most pronounced expression differences between VOC-exposed and control samples were observed at day 3 post-sowing (Figure 3), corresponding to early root radicle emergence and growth. No significant transcriptional alterations were detected at earlier time points (24 h and 48 h post-sowing), indicating that fungal VOCs trigger a rapid and early, yet transient, modulation of auxin and cytokinin pathways specifically during initial root establishment.

Among the auxin-related genes, the influx carrier *EIR1* showed the strongest response, with significantly higher transcript levels compared to non-exposed plants at 3 dps (Figure 3F). *AUX1* and *SHY2/IAA3* expression was also significantly up-regulated in the presence of fungal VOCs at this early root stage (Figure 3B, D). Regarding CK-related genes, *IPT3* showed no statistically significant differences upon exposure to fungal VOCs across any of the time points analyzed (Figure 3H), whereas *PUP14* expression was significantly down-regulated at 3 dps in the presence of fungal VOCs (Figure 3J).

Taken together, these results indicate that fungal VOCs primarily target auxin transport and signaling during early seedling establishment and initial root growth, while exerting a minimal transcriptional influence during earlier seed imbibition stages or on later CK biosynthesis and transport pathways.

### Fungal VOCs promote fresh weights and lateral root formation in Arabidopsis plants under control and osmotic stress conditions

To explore how the early molecular changes translate into root system development, we further analyzed primary root elongation and lateral root formation in CKs and auxin mutants under control (Supplementary Figure 5A) and osmotic stress conditions (Supplementary Figure 5B).

Fresh weights increased in the presence of fungal VOCs in Col-0 and in all the mutants analyzed, both under control and osmotic stress (Figures 4K-O and 5K-O), except for the *aux1-7* mutant under control conditions, where this increase was only observed in the presence of *S. indica* (Figure 4O).

**Figure 4.**
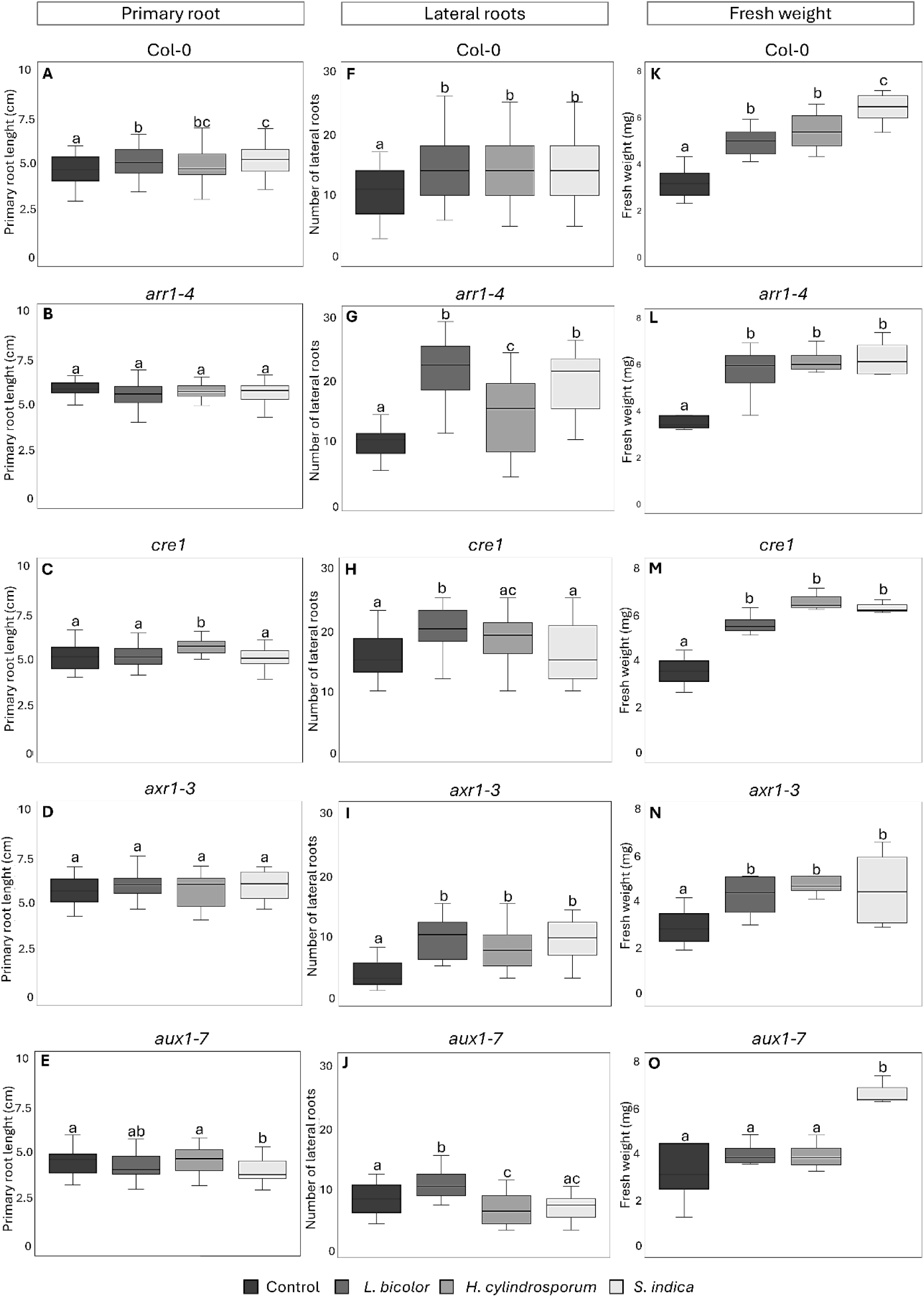
Analysis of 10-day-old seedlings of Col-0, *arr1-4*, *cre1*, *axr1-3*, and *aux1-7* in the absence and presence of fungal VOCs under control conditions. A-E. Primary root length (cm). F-J. Number of lateral roots. K-O. Total fresh weight (mg). Data are shown as boxplots of five biological replicates (n=5 plates) per genotype and condition; each plate contained 20 seeds per genotype. Different letters indicate statistical differences (p<0.05) among treatment means using one-way ANOVA and Tukey’s test.

**Figure 5.**
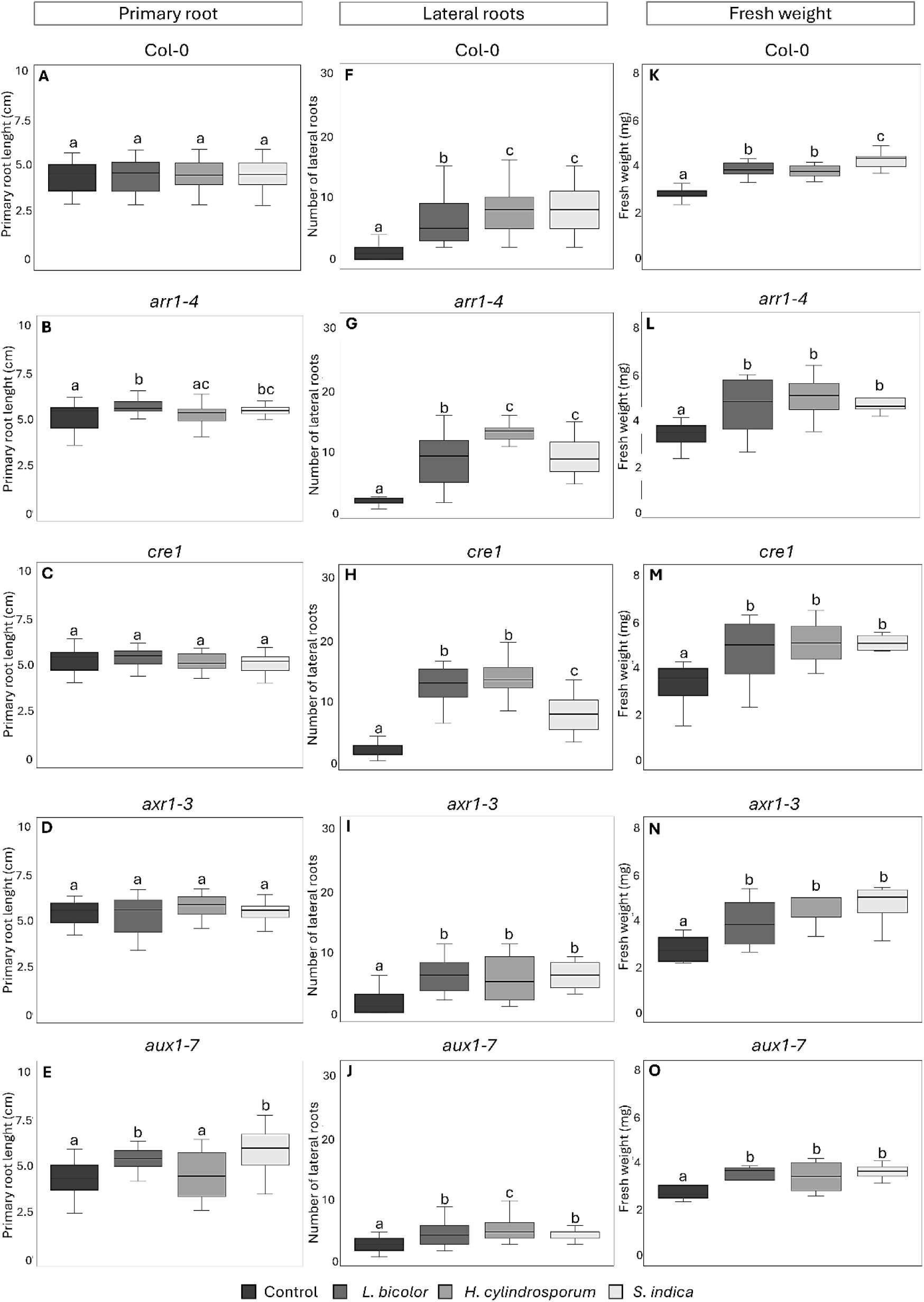
Analysis of 10-day-old seedlings of Col-0, *arr1-4*, *cre1*, *axr1-3*, and *aux1-7* in the absence and presence of fungal VOCs under osmotic stress conditions (sorbitol 100 mM). A-E. Primary root length (cm). F-J. Number of lateral roots. K-O. Total fresh weight (mg). Data are shown as boxplots of five biological replicates (n=5 plates) per genotype and condition; each plate contained 20 seeds per genotype. Different letters indicate statistical differences (p<0.05) among treatment means using one-way ANOVA and Tukey’s test.

The number of lateral roots increased in Col-0 seedlings and the mutant lines in the presence of all fungal VOCs, both under control and osmotic conditions (Figures 4F, 8F), except for *aux1-7* (Figure 4J) and *cre1* (Figure 4H) that only increased in the presence of VOCs emitted by *L. bicolor* (Figure 4H).

The length of the primary root of Col-0 seedlings increased in the presence of fungal VOCs (Figure 4A) under control conditions. In the CKs resistant mutant *cre1*, the length of the primary root increases only in the presence of fungal VOCs emitted by *H. cylindrosporum* (Figure 4C). However, in the auxin-resistant mutant *aux1-7*, the length of the primary root decreases in the presence of VOCs emitted by *S. indica* (Figure 4E). No further differences in primary root length were observed in the presence of VOCs emitted by any of the fungi (Figure 4B and 4D). On the other hand, under osmotic stress, primary root length significantly increased in the *arr1-4* mutant in the presence of fungal VOCs emitted by *L. bicolor* compared to the absence of these VOCs (Figure 5B). In the *aux1-7* mutant, an increase in primary root length was observed in the presence of VOCs emitted by both *L. bicolor* and *S. indica* (Figure 5E). No further significant differences in primary root length were observed in the presence of VOCs emitted by fungi in any of the other cases analyzed (Figure 5A, C, D).

Together, these data suggest that the phenotypic promotion triggered by fungal VOCs relies on a coordinated temporal balance also under stress conditions. Auxin transport operates as the primary trigger during early VOCs perception, whereas the classic auxin-cytokinin antagonism emerges at later developmental stages, with cytokinin signaling functioning as a homeostatic checkpoint to prevent overgrowth and balance root architecture.

### Fungal VOCs independently regulate glucosinolate and leucine metabolism, nitric oxide signaling and ion transport

To further analyze the extent to which fungal VOCs modulate molecular pathways related to the phenotypes observed at 10 days, we performed a root transcriptional profiling. Overall, fungal VOCs upregulated more genes than they repressed compared to non-exposed controls, both under control and osmotic stress conditions (Figure 6A and Supplementary Figure 6). VOCs emitted by *H. cylindrosporum* and *S. indica* induced a similar number of differentially expressed genes (DEGs) under both control (189/206 upregulated, 71/79 downregulated, respectively) and osmotic stress conditions (108/90 upregulated, 47/50 downregulated). In contrast, *L. bicolor* VOCs induced the fewest DEGs, with 34 upregulated and 4 downregulated genes under control conditions, and 23 upregulated and 12 downregulated under osmotic stress (Figure 6A and Supplementary Figure 6).

**Figure 6.**
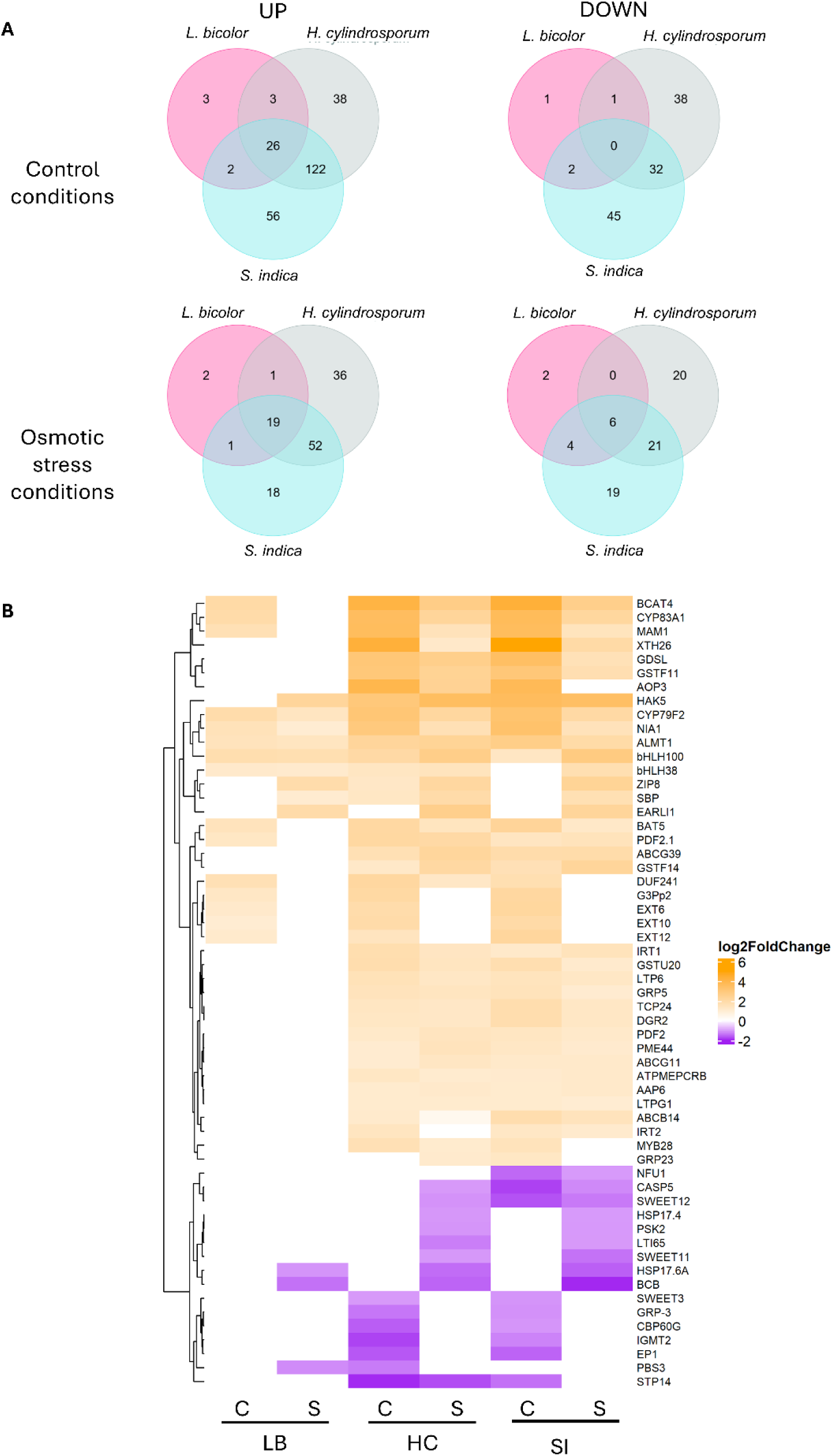
Root transcriptome analysis of Arabidopsis 10 days-old roots under control and osmotic stress conditions (100 mM sorbitol). A. Venn diagrams of differentially expressed genes (DEGs) exposed to VOCs emitted by *L. bicolor*, *H. cylindrosporu*, or *S. indica*, under non-stressed (top diagrams) or osmotic stress conditions (bottom diagrams), comparing in all cases with plants not exposed to fungal VOCs. Upregulated genes are shown as UP and downregulated genes as DOWN. B. DEGs analysis of genes induced (orange) and repressed (purple) under control (C) and osmotic stress (S) conditions. The Y-axis on the right shows the names of the different genes represented. Data are from three biological replicates (n=3) per condition and DEGs were identified with DESeq2 (|log₂(fold change)| ≥ 1.0, q-value ≤ 0.05).

Gene ontology (GO) enrichment analysis (Supplementary Figures 7–9) revealed that processes involved in the biosynthesis and metabolism of S-glucosides, glucosinolates, and leucine were significantly enhanced by all fungal VOCs under both control and osmotic stress conditions (Supplementary Figures 8 and 9). Consistently, key upregulated genes driving these metabolic GO terms included *BCAT4* (Schuster et al., 2006) (Figure 6B, Supplementary Figures 10A and 10B), *BAT5* (Gigolashvili et al., 2009), and *MAM1* (Kroymann et al., 2003) (Figure 6B). Additionally, genes involved in biotic/abiotic stress responses, such as *GDSL*, previously associated with symbiotic interactions with *S. indica* (Vahabi et al., 2015; 2013), were induced (Figure 6B).

GO terms associated with intracellular ion homeostasis and cell transporters were enriched under control conditions in response to *H. cylindrosporum* (Supplementary Figure 7A) and *S. indica* VOCs (Supplementary Figure 9), as well as under osmotic stress with the VOCs of the tree fungi (Supplementary Figure 8B). This was further supported by the induction of specific ion transporters and iron-deficiency regulators, including the iron transporter *IRT1* (Vert et al., 2021), the transcription factors *bHLH38* and *bHLH100* (Gao & Dubos, 2024), the aluminum transporter *ALMT1* (Wang et al., 2021), and the potassium transporter *HAK5* (Hong et al., 2013) (Figure 6B, Supplementary Figures 10C and 10D). Conversely, sugar transporters such as *STP14* (Poschet et al., 2010) and *SWEET12* (Le Hir et al., 2015; Chen et al., 2012) were consistently repressed by *H. cylindrosporum* and *S. indica* VOCs regardless of stress conditions (Figure 6B, Supplementary Figures 10E and 10F).

Fungal VOCs also modulated hormonal signaling and cell wall reorganization. Within hormone-related pathways, auxin-associated genes *ABCB14* (Cho & Cho, 2013) and *AGL14* (Garay-Arroyo et al., 2013) were upregulated by *H. cylindrosporum* and *S. indica* VOCs under both control and osmotic stress conditions, whereas *PIN5* (Mravec et al., 2009) was specifically induced by *L. bicolor* VOCs under control conditions (Figure 6B). Furthermore, the nitric oxide pathway gene *NIA1* was induced across all analyzed conditions (Vazquez et al., 2019; Hao et al., 2010) (Figure 6B). Lastly, genes linked to cellular reorganization and root differentiation, such as xyloglucan synthesis-related *XTH26* (Becnel et al., 2006), were also upregulated under osmotic stress conditions in the presence of VOCs from all three fungi.

### Fungal VOCs regulate potassium and iron transporters

To validate key RNA-seq findings and explore their functional relevance, we analyzed mutants and gene expression pathways associated with ion transport and nitric oxide (NO) biosynthesis. Given that *HAK5* was significantly induced by fungal VOCs under control and osmotic stress treatments (Figure 6B, Supplementary Figures 10C and 10D) and considering the well-established role of potassium transporters and channels in modulating root system architecture (Li et al., 2017; Brauer et al., 2016), we next evaluated the functional relevance of this pathway. We analyzed the root phenotypes of *hak5-2* and *akt1-2 Arabidopsis* mutants under control conditions (Figure 7A), which represent interconnected core modules critical for auxin (Chatterjee et al., 2022) and ethylene-mediated (Jung et al., 2009) root development. Regarding primary root length, *akt1-2* seedlings showed an increase in response to *H. cylindrosporum* and *S. indica* VOCs but a decrease with *L. bicolor* VOCs, whereas *hak5-2* exhibited increased length with *S. indica* but a reduction with *H. cylindrosporum* (Figure 7B). As for lateral roots, although both mutant lines displayed increases in lateral root number upon exposure to fungal VOCs, this induction was less pronounced than that observed in Col-0 (Figure 4F and Figure 7C). Finally, fresh weight did not increase in either *akt1-2* or *hak5-2* mutants in response to any of the tested fungal VOCs (Figure 7D).

**Figure 7.**
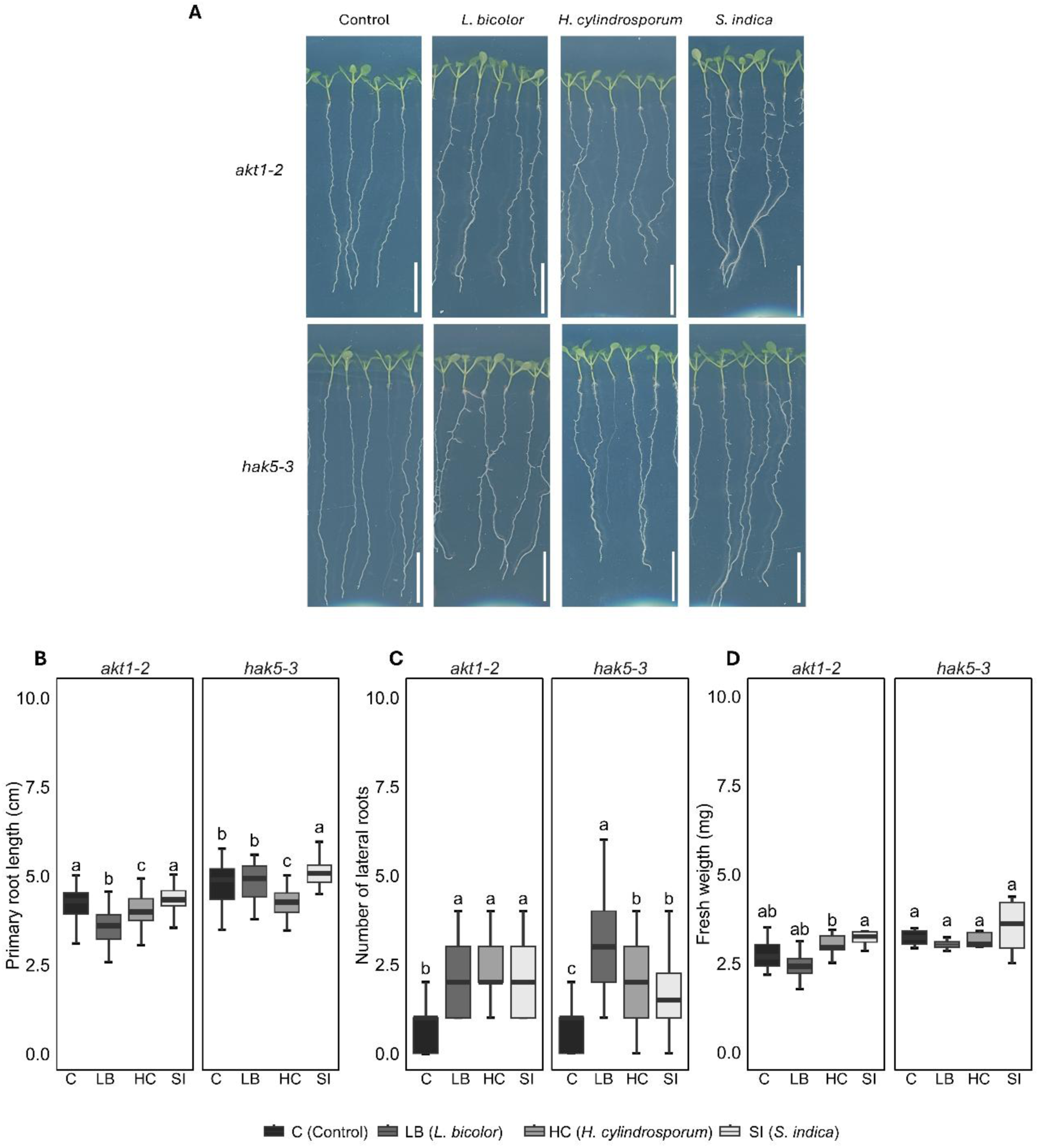
Analysis of 10 day-old seedling development of *HAK5*-related potassium transporter mutants (*hak5-2* and *akt1-2*) in the absence and presence of fungal VOCs under control conditions. A. Phenotype of *akt1-2* and *hak5-2*. Scale bar = 1 cm. B. Primary root length (cm). C. Number of lateral roots. D. Total fresh weight (mg). Data are shown as boxplots of five biological replicates (n=5 plates) per genotype and condition; each plate contained 20 seeds per genotype. Different letters indicate statistical differences (p<0.05) among treatment means using one-way ANOVA and Tukey’s test.

Regarding NO signaling, the *nia1;nia2* double mutant exhibited reduced, but still significant, increases in fresh weight and lateral root number under VOC exposure from all three fungi (Figures 8A–C). In contrast, the *noa1-2;nia1;nia2* triple mutant completely abolished VOC-induced increases in both parameters (Figures 8E–G). Because NO acts as a key signaling molecule regulating iron-dependent genes and their physiological uptake and mobilization processes (Tewari et al., 2021; Buet & Simontacchi, 2015), the expression of *IRT1* was further analyzed by RT-qPCR. *IRT1* relative expression levels were significantly reduced in both *nia1;nia2* double and *noa1-2;nia1;nia2* triple mutants under fungal VOC exposure (Figures 8D and 8H), with a particularly strong repression observed in the triple mutant exposed to *L. bicolor* and *H. cylindrosporum* VOCs (Figure 8H).

**Figure 8.**
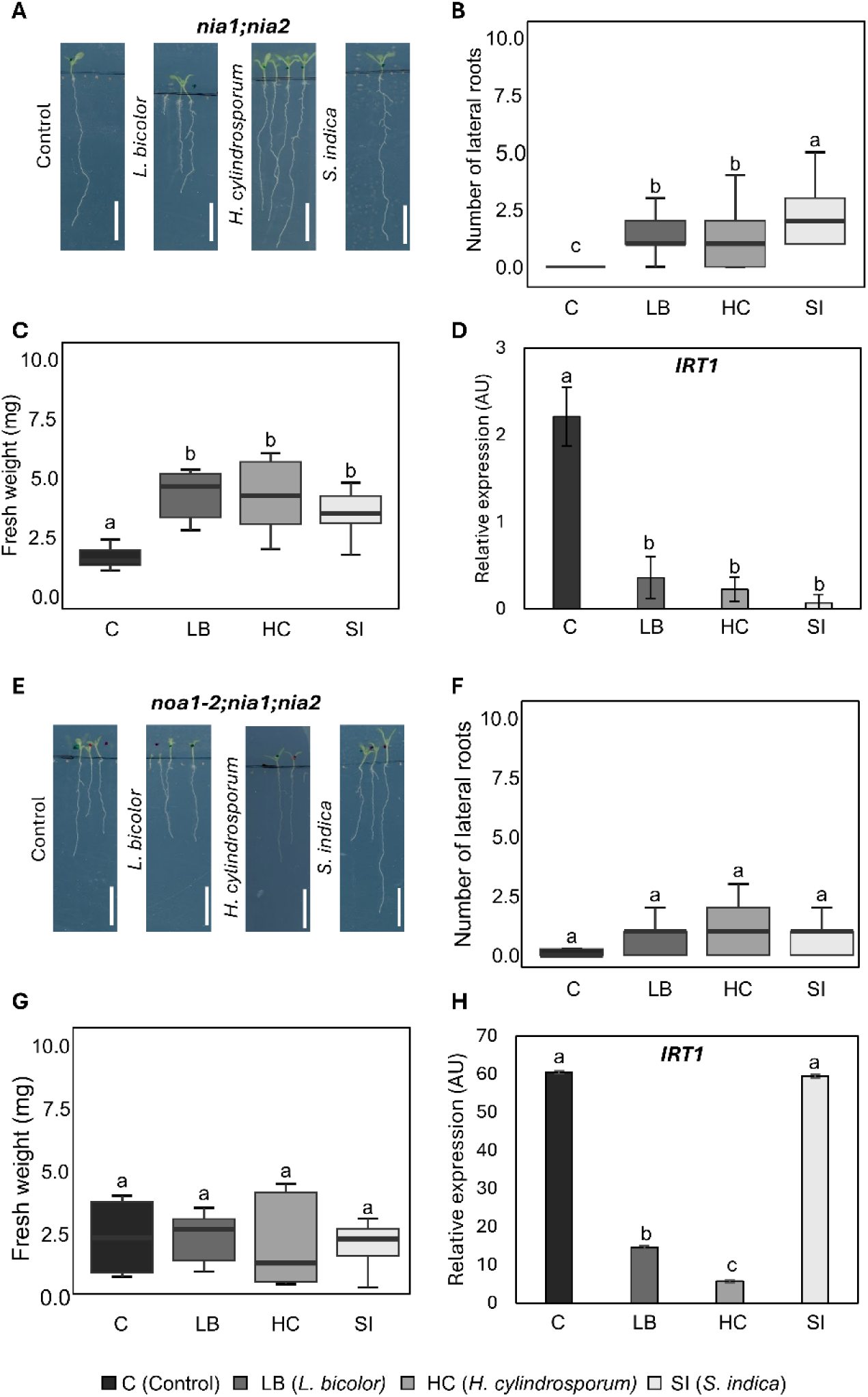
Analysis of 14 day-old seedling development of *nia1;nia2* and *noa1-2;nia1;nia2* NO-deficient mutants in the absence and presence of fungal VOCs under control conditions. A. Phenotype of *nia1;nia2*. Scale bar = 1 cm. B. Number of lateral roots of *nia1;nia2*. C. Total fresh weight of *nia1;nia2* expressed in mg. D. Relative expression of *IRT1* in *nia1;nia2* mutant. E. Phenotype of *noa1-2;nia1;nia2*. Scale bar= 1 cm. F. Number of lateral roots of *noa1-2;nia1;nia2*. G. Total fresh weight of *noa1-2;nia1;nia2* expressed in mg. H. Relative expression of *IRT1* in *noa1-2;nia1;nia2* mutant. Data are shown as boxplots of five biological replicates (n=5 plates) per genotype and condition; each plate contained 20 seeds per genotype. Different letters indicate statistical differences (p<0.05) among treatment means using one-way ANOVA and Tukey’s test.

Altogether, our data indicates that fungal VOCs trigger a multi-pronged response involving stress tolerance metabolism, nitric oxide signaling, and ion homeostasis. These pathways operate as parallel functional modules that translate VOC perception into enhanced root development and stress tolerance.

## Discussion

### The effect of fungal VOCs on plant development depends on a consortium of compounds

The volatilomes of the studied fungi exhibited high diversity and distinct chemical profiles (Figures 1A and 1C), while sharing specific core compounds such as 2-methyl-1-propanol and 3-methyl-1-butanone (Hung et al., 2015; Schnürer et al., 1999). Exposure to these fungal VOCs enhanced plant growth, leaf chlorophyll content, and lateral root formation (Figures 2C, 4F–J, 5F–J, and Supplementary Figure 5), aligning with well-documented benefits reported for various mycorrhizal (Casarrubia et al., 2016; Ditengou et al., 2015; Wu et al., 2013), *Trichoderma* spp. (Garnica-Vergara et al., 2016; Uddin et al., 2018), and pathogenic fungi (Casarrubia et al., 2016; Moisan et al., 2021). Notably, although individual candidate VOCs (2-methylpropanal, 3-methyl-1-butanone, 3-octanone, octanal, and farnesylacetone) partially increased fresh weight and lateral roots, none fully replicated the robust phenotypic effects induced by the complete natural volatile profile (Figures 3, 4, and Supplementary Figure 2A). Consistent with previous findings in *Serendipita* spp. (Venneman et al., 2020), these results indicate that isolated compounds are not, on their own, the sole drivers of these developmental responses, suggesting that synergistic interactions within complex volatile mixtures or additional unidentified compounds underline the full phenotypic effects observed. Future studies testing binary or ternary mixtures of these VOCs could help elucidate whether synergistic interactions among specific compound combinations underline the full developmental effects observed with fungal volatile profile.

### Fungal VOCs induction of lateral roots are controlled by the temporal auxin/cytokinin balance

Auxins and CKs are antagonistic hormonal pathways in root development. Their interaction plays a crucial role in the differentiation of root stem cells during early embryogenesis, as well as in the necessity for CKs to antagonize auxin flow for lateral root initiation and development (Kurepa & Smalle, 2022; Kurepa et al., 2019; Perilli et al., 2010). Our time-course results showed that fungal VOCs modulates auxin transport (*AUX1 and EIR1)* and signaling (*SHY2/IAA3*) genes at early developmental steps (3 days) (Figure 3A-F), with rapid but transient expression changes, while CK pathways show minimal modulation under control conditions (Figure 3G-J). This specific activation of auxin machinery is highly consistent with the establishment of an auxin gradient essential for lateral root founder cell priming within the first 24–72 hours (Bielach et al., 2012; De Smet et al., 2007) of VOCs perception. Notably, the promotion of lateral root proliferation by *L. bicolor* VOCs in *Populus tremula × alba* has been previously linked to the induction of the auxin pathway through the alteration of PIN-mediated polar auxin transport (Ditengou et al., 2015). Our data strongly support this mechanism across multiple fungal species, as evidenced by the early upregulation of *AUX1* and particularly *EIR1* (Figure 3), which encodes the PIN-family efflux carrier PIN2 (Luschnig et al., 1998).

Consequently, at later developmental stages (10 days), this early molecular switch translates into long-term root architecture remodeling, where the functional integration of the auxin/CKs balance becomes phenotypically apparent. Both under control and osmotic stress conditions, an increase in lateral root number was more pronounced in CKs-resistant mutants and much less significant in auxin-resistant mutants (Figure 4 and 5). In auxin mutants (Figures 4G, H, Figure 5G, H), a smaller increase in lateral root number compared to Col-0 was observed, as auxins failed to exert their promoting effect on lateral roots (Jing & Strader, 2019; Fukaki & Tasaka, 2009; Casimiro et al., 2003). Conversely, CK-resistant mutants (Figures 3I, J, Figure 4I, J) exhibited a higher increase in lateral roots than Col-0, indicating that the absence of CK perception induced the effect of the auxins, thereby resulting in more lateral roots. This phenotypic response supports the observations by Venneman et al. (2020), that described the lateral root proliferation triggered by *S. indica* VOCs occurs through a modulation of both auxin and CKs pathways.

Finally, our transcriptomic results at 10 days-old seedling provides comprehensive molecular evidence of how this long-term root remodeling is sustained. These results demonstrate that fungal VOCs emitted by *L. bicolor*, *H. cylindrosporum*, and *S. indica* actively maintain the induction of alternative auxin transport and signaling modules under both control and osmotic stress conditions (Figure 6). Beyond the initial induction of *AUX1* (Goh et al., 2012), fungal VOCs exposure upregulated key players involved in intracellular auxin homeostasis and distribution, such as the transporter *PIN5* (Mravec et al., 2009), the auxin/proton symporter *ABCB14* (Cho & Cho, 2013), and the transcription factor *AGL14*, a well-known positive regulator of lateral root development via auxin transport modulation (Garay-Arroyo et al., 2013). Taken together, this multi-layered genetic and transcriptional regulation highlights a model where fungal VOCs trigger an early, transient auxin transport regulation that sequentially bypasses the antagonism between cytokinin and auxin pathways, establishing a robust, auxin-driven developmental program to optimize root system development.

### Fungal VOCs induce abiotic stress tolerance in *Arabidopsis thaliana* plants

The positive effects of fungal VOCs on the improvement of the tolerance of against abiotic stress has been previously described in other fungi-plant systems as *S. indica* in *Ocinum basilicum* (Fraj & Werbrouck, 2023)*, Tolypocladium inflatum* in tomato (Chiang et al., 2024) and bacteria like *Pseudomonas spp.* in *Mentha piperita* (Gil et al., 2023). Importantly, ectomycorrhizal fungi like *L. bicolor* and *H. cylindrosporum* cannot physically colonize Brassicaceae species such as *Arabidopsis thaliana* (Smith & Read, 2010), in stark contrast to the endophyte *S. indica*, which establishes a functional mutualistic interaction (González Ortega-Villaizán et al., 2024; Opitz et al., 2021). Despite these contrasting lifestyles and host specificities, all three fungi induced strong developmental and stress-tolerance responses in *A. thaliana* through their volatile emissions. Our root transcriptome analyses highlight a coordinated response involving interconnected pathways that enhance the stress tolerance system of *A. thaliana*, demonstrating that volatile-mediated plant growth promotion operates independently of physical root colonization.

Within this network, the upregulation of leucine and glucosinolate biosynthesis pathways represents a core VOC-responsive metabolic module. Although leucine biosynthesis and leucine-rich repeat proteins have been shown to be essential for salt stress tolerance in Arabidopsis seedlings (Zhao et al., 2018), and leucine-related signaling can modulate primary root elongation and cell wall integrity (Ravelo-Ortega et al., 2021; Van der Does et al., 2017; Draeger et al., 2015), their induction as a direct response to fungal volatile exposure has not been previously reported. Our findings reveal that fungal VOCs trigger these leucine metabolism genes (Figure 9B), actively contributing to the observed root architecture remodeling.

This metabolic shift is reinforced by the concurrent upregulation of glucosinolate biosynthesis pathways (Figure 6B), which are critical for chemical defense and osmotic adjustments in Brassicaceae (Burow & Halkier, 2017; Petersen et al., 2002). Notably, the differentially expressed genes induced by fungal VOCs under control conditions largely overlap with those altered under osmotic stress (Figure 6). This comprehensive metabolic and transcriptional reprogramming by glucosinolates and leucine pathways likely establishes a priming state, enabling plants to deploy faster and more effective stress responses upon a subsequent environmental challenge (Aswathi et al., 2025; Jung et al., 2012). Ultimately, this VOC-mediated priming manifests as early root architecture adaptations that enhance water and nutrient uptake capacity prior to stress onset (Laller et al., 2023; Yasmin et al., 2021).

### Fungal VOCs regulate ion and sugar transporters and nitric oxide under osmotic stress

Arabidopsis seedlings exposed to fungal VOCs induced genes related to ion and sugar transport both under control and osmotic stress conditions (Figure 6B), with special interest on HAK5, a high-affinity potassium transporter (Hong et al., 2013), and IRT1, an iron transporter (Vert et al., 2021), known for their function in plant stress responses.

Given that HAK5 is required for broad abiotic stress responses (Brauer et al., 2016), its upregulation by fungal VOCs under both control and osmotic stress conditions (Figure 6B) identifies it as a key node in the regulation of volatile-mediated signaling in plants. We corroborate this signaling role, since the *hak5-2* and *akt1-2* mutants exhibited significantly attenuated lateral root proliferation compared to Col-0 and a complete abolishment of fresh weight increases upon fungal VOC exposure (Figure 7). Rather than operating strictly as nutrient acquisition channels under nutritional deficiency, HAK5 and AKT1 transporters are increasingly recognized as core biophysical components that modulate root system architecture by regulating cellular membrane potential, turgor pressure, and elongation (Chatterjee et al., 2022; Li et al., 2017). The phenotype observed in *hak5-2* and *akt1-2* exposed to fungal VOCs can be explained by the intricate crosstalk between transporters and hormone signaling pathways during root development. Specifically, AKT1 has been directly implicated in modulating auxin-mediated root growth responses (Li et al., 2017), while the transcriptional regulation of HAK5 is tightly controlled by auxin response factors, such as ARF2, which act as molecular switches coupling hormone signaling to transporter expression (Zhao et al., 2016). Furthermore, ethylene signaling acts as a critical upstream mediator regulating plant growth adjustments and abiotic stress tolerance linked to HAK5 transport activity (Jung et al., 2009). Consequently, these results indicate that functional *HAK5* and *AKT1* modules are stringently required to transduce VOC-induced auxin and ethylene signals into physical lateral root development and biomass gain.

The transcriptional induction of *IRT1* (Figure 6B) is known to trigger root architecture remodeling under iron deficiency, specifically increasing root hair density and lateral root formation (Vert et al., 2021; Connorton et al., 2017). Also, transcriptomic data revealed the upregulation of *NIA1* in the presence of three fungal VOCs under both control and osmotic stress conditions (Figure 6B). *NIA1* encodes a nitrate reductase isoform that reduces nitrate to nitric oxide (NO) (Olas & Wahl, 2019), a critical gaseous signaling molecule that regulates iron-responsive genes and coordinates physiological iron uptake and mobilization (Tewari et al., 2021; Buet & Simontacchi, 2015). Crucially, *nia1;nia2* double mutants and *noa1-2;nia1;nia2* triple mutants exhibited a severe reduction in *IRT1* expression upon VOC exposure, particularly upon exposure to *L. bicolor* and *H. cylindrosporum* VOCs (Figure 8). Consequently, these mutants failed to exhibit VOC-induced increases in fresh weight and lateral root number (Figure 8). Since NO-deficient mutants not only possess impaired NO production but also display general disruptions in metal homeostasis, such as iron (Zhu et al., 2015), our findings strongly demonstrate that NO acts as a critical signaling hub linking VOC perception to iron homeostasis and root development improvement.

Additionally, the sugar transporters *STP14* and *SWEET12* were significantly repressed in our transcriptomic data (Figure 6B). The regulation of these carriers is known to be key in the response to various types of abiotic stress, including drought (Du et al., 2022) and biotic interactions (Breia et al., 2021). These findings reinforce the central role of transporters in VOC-mediated signaling, potentially triggering downstream molecular processes inside root cells that are yet to be fully elucidated.

Taken together, our results indicate that fungal VOCs enhance plant stress tolerance through coordinated molecular reprogramming of auxin transport, ion homeostasis, and defense metabolism. VOCs trigger an early, rapid modulation of auxin transporters (*AUX1, PIN2, SHY2*) within the first 3 days, followed by a subsequent requirement to bypass the cytokinin antagonistic brake at 10 days, thereby establishing the precise hormonal balance essential for lateral root proliferation under both control and osmotic stress conditions. Transcriptomic analyses revealed robust activation of glucosinolate/leucine biosynthetic pathways and ion transporters (*HAK5, IRT1, ALMT1*), where functional validation confirms nitric oxide biosynthesis (*NIA1*) and potassium transport networks as critical signaling hubs linking VOC perception to enhanced root system architecture. Beyond developmental regulation, fungal VOCs establish a transcriptional priming state that confers cross-tolerance to osmotic stress through preemptive defense activation, independent of physical fungal contact. While fungal inoculation improves crop performance under stress (Zeng et al., 2025; Adedayo & Babalola, 2023), it risks disrupting soil microbiota (O’Callaghan et al., 2022); VOC-based strategies offer a targeted alternative, harnessing microbial signaling benefits without ecological drawbacks, thereby offering significant potential for sustainable agriculture and forestry applications under diverse abiotic stresses.

## Supplementary Data

**Supplementary Table 1.** Oligonucleotides used for qPCR, including sequences and melting temperatures.

**Supplementary Figure 1.** Schematic representation of the experimental set-ups used for fungal VOC application.

**Supplementary Figure 2.** Volatile compound profiles in the growth medium after 6 days of fungal VOC exposure, and phenotypic analysis of Col-0 seedlings treated with selected individual VOCs.

**Supplementary Figure 3.** Growth phenotypes of abscisic acid signaling mutants exposed to VOCs from *L. bicolor* and *S. indica*.

**Supplementary Figure 4.** Growth phenotypes of ethylene, gibberellin, and jasmonic acid signaling mutants exposed to VOCs from *L. bicolor* and *S. indica*.

**Supplementary Figure 5.** Overall seedling phenotypes of 10-day-old *Arabidopsis* plants exposed to VOCs from *L. bicolor*, *H. cylindrosporum*, and *S. indica*.

**Supplementary Figure 6.** Volcano plots showing differentially expressed genes in 10-day-old Col-0 roots exposed to fungal VOCs under control and osmotic stress conditions.

**Supplementary Figures 7–9.** Gene Ontology enrichment analyses of differentially expressed genes in roots exposed to VOCs from *L. bicolor*, *H. cylindrosporum*, and *S. indica*, respectively.

**Supplementary Figure 10.** qPCR validation of selected genes from the transcriptomic profiling of roots exposed to fungal VOCs under control and osmotic stress.

## Acknowledgements

We thank the Elemental Analysis, Chromatography, and Mass Spectrometry Facility (NUCLEUS, University of Salamanca) for technical support and access to analytical instrumentation for VOC metabolite profiling in agar samples. We also acknowledge the Molecular Biology Service (ACTI, University of Murcia) for RNA-seq library preparation, sequencing, and associated quality control.

## Author contributions

MC-P and EM conceived and designed the experiments; EM performed the experiment and sample measurements; EM, OL and MC-P analyzed the data and JF-D, SR-S and PM-M analyzed the volatiles profile; EM, MC-P and OL drafted the manuscript, with contributions from all coauthors.

## Data availibility

The data that supports the findings within the study are available from the corresponding authors upon request. The sequence data can be found at the Gene Expression Omnibus (GEO) database (NCBI) under the following accession number GSE308215.

## References

Adedayo, A. A., & Babalola, O. O. 2023. Fungi That Promote Plant Growth in the Rhizosphere Boost Crop Growth. Journal of Fungi, 9(2), 239. doi: 10.3390/jof9020239

Arabidopsis Genome Initiative. 2000. Analysis of the genome sequence of the flowering plant Arabidopsis thaliana. Nature, 408(6814), 796–815. doi: 10.1038/35048692

Aswathi, K. P. R., Ul-Allah, S., Puthur, J. T., Siddique, K. H. M., Frei, M., & Farooq, M. 2025. The Plant Mind: Unraveling Abiotic Stress Priming, Memory, and Adaptation. Physiologia Plantarum, 177(4). doi: 10.1111/ppl.70372

Becnel, J., Natarajan, M., Kipp, A., & Braam, J. 2006. Developmental Expression Patterns of Arabidopsis XTH Genes Reported by Transgenes and Genevestigator. Plant Molecular Biology, 61(3), 451–467. doi: 10.1007/s11103-006-0021-z

Berardini, T. Z., Reiser, L., Li, D., Mezheritsky, Y., Muller, R., Strait, E., & Huala, E. 2015. The arabidopsis information resource: Making and mining the “gold standard” annotated reference plant genome. Genesis, 53(8), 474–485. doi: 10.1002/dvg.22877

Bertaux, J., Schmid, M., Prevost-Boure, N. C., Churin, J. L., Hartmann, A., Garbaye, J., & Frey-Klett, P. 2003. In Situ Identification of Intracellular Bacteria Related to *Paenibacillus* spp. in the Mycelium of the Ectomycorrhizal Fungus *Laccaria bicolor* S238N. Applied and Environmental Microbiology, 69(7), 4243–4248. doi: 10.1128/AEM.69.7.4243-4248.2003

Bielach, A., Podlešáková, K., Marhavý, P., Duclercq, J., Cuesta, C., Müller, B., Grunewald, W., Tarkowski, P., & Benkováa, E. 2012. Spatiotemporal Regulation of Lateral Root Organogenesis in Arabidopsis by Cytokinin. The Plant Cell, 24(10), 3967–3981. doi: 10.1105/TPC.112.103044

Brauer, E. K., Ahsan, N., Dale, R., Kato, N., Coluccio, A. E., Piñeros, M. A., Kochian, L. v., Thelen, J. J., & Popescu, S. C. 2016. The Raf-like Kinase ILK1 and the High Affinity K+ Transporter HAK5 Are Required for Innate Immunity and Abiotic Stress Response. Plant Physiology, 171(2), 1470–1484. doi: 10.1104/PP.16.00035

Breia, R., Conde, A., Badim, H., Fortes, A. M., Gerós, H., & Granell, A. 2021. Plant SWEETs: from sugar transport to plant–pathogen interaction and more unexpected physiological roles. Plant Physiology, 186(2), 836–852. doi: 10.1093/plphys/kiab127

Buet, A., & Simontacchi, M. 2015. Nitric oxide and plant iron homeostasis. Annals of the New York Academy of Sciences, 1340(1), 39–46. doi: 10.1111/nyas.12644

Burow, M., & Halkier, B. A. 2017. How does a plant orchestrate defense in time and space? Using glucosinolates in Arabidopsis as case study. Current Opinion in Plant Biology, 38, 142–147. doi: 10.1016/j.pbi.2017.04.009

Calvo-Polanco, M., Armada, E., Zamarreño, A. M., García-Mina, J. M., & Aroca, R. 2019. Local root ABA/cytokinin status and aquaporins regulate poplar responses to mild drought stress independently of the ectomycorrhizal fungus *Laccaria bicolor*. Journal of Experimental Botany, 70(21), 6437–6446. doi: 10.1093/JXB/ERZ389

Câmara, J. S., Perestrelo, R., Ferreira, R., Berenguer, C. v., Pereira, J. A. M., & Castilho, P. C. 2024. Plant-Derived Terpenoids: A Plethora of Bioactive Compounds with Several Health Functions and Industrial Applications—A Comprehensive Overview. Molecules, 29(16), 3861. doi: 10.3390/molecules29163861

Carlson M. 2024. _org.At.tair.db: Genome wide annotation for Arabidopsis_. R package version 3.19.1. (n.d.).

Casarrubia, S., Sapienza, S., Fritz, H., Daghino, S., Rosenkranz, M., Schnitzler, J.-P., Martin, F., Perotto, S., & Martino, E. 2016. Ecologically Different Fungi Affect Arabidopsis Development: Contribution of Soluble and Volatile Compounds. PLOS ONE, 11(12), e0168236. doi: 10.1371/journal.pone.0168236

Casimiro, I., Beeckman, T., Graham, N., Bhalerao, R., Zhang, H., Casero, P., Sandberg, G., & Bennett, M. J. 2003. Dissecting Arabidopsis lateral root development. Trends in Plant Science, 8(4), 165–171. doi: 10.1016/S1360-1385(03)00051-7

Chatterjee, M., Ghosh, P., Sen, S., Sinha, D., & Ganguly, S. 2022. The Function of HAK as K+ Transporter and AKT as Inward-Rectifying Agent in the K+ Channel. Plant Metal and Metalloid Transporters, 227–243. doi: 10.1007/978-981-19-6103-8_11

Chen, L.-Q., Qu, X.-Q., Hou, B.-H., Sosso, D., Osorio, S., Fernie, A. R., & Frommer, W. B. 2012. Sucrose Efflux Mediated by SWEET Proteins as a Key Step for Phloem Transport. Science, 335(6065), 207–211. doi: 10.1126/science.1213351

Chen, T., & Fluhr, R. 2018. Singlet Oxygen Plays an Essential Role in the Root’s Response to Osmotic Stress. Plant Physiology, 177(4), 1717–1727. doi: 10.1104/pp.18.00634

Chiang, C.-Y., Chang, C.-H., Tseng, T.-Y., Nguyen, V.-A. T., Su, P.-Y., Truong, T.-T. T., Chen, J.-Y., Huang, C.-C., & Huang, H.-J. 2024. Volatile Compounds Emitted by Plant Growth–Promoting Fungus *Tolypocladium inflatum* GT22 Alleviate Copper and Pathogen Stress. Plant And Cell Physiology, 65(2), 199–215. doi: 10.1093/pcp/pcad120

Chiu, C. H., Roszak, P., Orvošová, M., & Paszkowski, U. 2022. Arbuscular mycorrhizal fungi induce lateral root development in angiosperms via a conserved set of MAMP receptors. Current Biology, 32(20), 4428–4437.e3. doi: 10.1016/j.cub.2022.08.069

Cho, M., & Cho, H. T. 2013. The function of ABCB transporters in auxin transport. Plant Signaling & Behavior, 8(2). doi: 10.4161/psb.22990

Connorton, J. M., Balk, J., & Rodríguez-Celma, J. 2017. Iron homeostasis in plants – a brief overview. Metallomics, 9(7), 813–823. doi: 10.1039/C7MT00136C

de Smet, I., Tetsumura, T., de Rybel, B., et al. 2007. Auxin-dependent regulation of lateral root positioning in the basal meristem of Arabidopsis. Development, 134(4), 681–690. doi: 10.1242/DEV.02753

di Battista, C., Selosse, M.-A., Bouchard, D., Stenström, E., & le Tacon, F. 1996. Variations in symbiotic efficiency, phenotypic characters and ploidy level among different isolates of the ectomycorrhizal basidiomycete *Laccaria bicolor* strain S 238. Mycological Research, 100(11), 1315–1324. doi: 10.1016/S0953-7562(96)80058-X

Ditengou, F. A., Müller, A., Rosenkranz, M., Felten, J., Lasok, H., van Doorn, M. M., Legué, V., Palme, K., Schnitzler, J.-P., & Polle, A. 2015. Volatile signalling by sesquiterpenes from ectomycorrhizal fungi reprogrammes root architecture. Nature Communications, 6(1), 6279. doi: 10.1038/ncomms7279

Dotson, B. R., Verschut, V., Flärdh, K., Becher, P. G., & Rasmusson, A. G. 2020. *The* Streptomyces *volatile 3-octanone alters auxin/cytokinin and growth in* Arabidopsis thaliana *via the gene family* KISS ME DEADLY. BioRxiv doi: 10.1101/2020.02.15.949685. [Preprint].

Draeger, C., Ndinyanka Fabrice, T., Gineau, et al. 2015. Arabidopsis leucine-rich repeat extensin (LRX) proteins modify cell wall composition and influence plant growth. BMC Plant Biology, 15(1), 155. doi: 10.1186/s12870-015-0548-8

Du, Y., Li, W., Geng, J., et al. 2022. Genome-wide identification of the SWEET gene family in *Phaseolus vulgaris* L. and their patterns of expression under abiotic stress. Journal of Plant Interactions, 17(1), 390–403. doi: 10.1080/17429145.2022.2044079

El Jaddaoui, I., Rangel, D. E. N., & Bennett, J. W. 2023. Fungal volatiles have physiological properties. Fungal Biology, 127(7–8), 1231–1240. doi: 10.1016/j.funbio.2023.03.005

El-Showk, S., Ruonala, R., & Helariutta, Y. 2013. Crossing paths: cytokinin signalling and crosstalk. Development, 140(7), 1373–1383. doi: 10.1242/dev.086371

Fraj, H., & Werbrouck, S. P. O. 2023. Constant and Intermittent Contact with the Volatile Organic Compounds of *Serendipita indica* Alleviate Salt Stress *In Vitro Ocimum basilicum* L. Applied Sciences, 13(3), 1776. doi: 10.3390/app13031776

Fresno, D. H., Solé-Corbatón, H., & Munné-Bosch, S. 2023. Water stress protection by the arbuscular mycorrhizal fungus *Rhizoglomus irregulare* involves physiological and hormonal responses in an organ-specific manner. Physiologia Plantarum, 175(1), e13854. doi: 10.1111/PPL.13854

Fukaki, H., & Tasaka, M. 2009. Hormone interactions during lateral root formation. Plant Molecular Biology, 69(4), 437–449. doi: 10.1007/s11103-008-9417-2

Gao, F., & Dubos, C. 2024. The Arabidopsis bHLH transcription factor family. Trends in Plant Science, 29(6), 668–680. doi: 10.1016/j.tplants.2023.11.022

Gao, C. H., Yu, G., & Cai, P. (2021). ggVennDiagram: An Intuitive, Easy-to-Use, and Highly Customizable R Package to Generate Venn Diagram. Frontiers in Genetics, 12, 706907. doi: 10.3389/FGENE.2021.706907

Garay-Arroyo, A., Ortiz-Moreno, E., de la Paz Sánchez, M., et al. 2013. The MADS transcription factor XAL2/AGL14 modulates auxin transport during Arabidopsis root development by regulating PIN expression. The EMBO Journal, 32(21), 2884–2895. doi: 10.1038/emboj.2013.216

Garnica-Vergara, A., Barrera-Ortiz, S., Muñoz-Parra, E., Raya-González, J., Méndez-Bravo, A., Macías-Rodríguez, L., Ruiz-Herrera, L. F., & López-Bucio, J. 2016. The volatile 6-pentyl-2 *H*-pyran-2-one from *Trichoderma atroviride* regulates *Arabidopsis thaliana* root morphogenesis via auxin signaling and *ETHYLENE INSENSITIVE 2* functioning. New Phytologist, 209(4), 1496–1512. doi: 10.1111/nph.13725

Ghaffari Jabbari, S., Fermoso Domínguez, J., Rodríguez Sufuentes, S., Nyberg, S. O., Sandnes Vehus, T., & Kofoed Nielsen, H. 2025. VOC emissions from commercial wood panels using PTR-MS for indoor air quality evaluation. Frontiers in Built Environment, 11, 1591669. doi: 10.3389/FBUIL.2025.1591669

Gigolashvili, T., Yatusevich, R., Rollwitz, I., Humphry, M., Gershenzon, J., & Flügge, U. I. 2009. The Plastidic Bile Acid Transporter 5 Is Required for the Biosynthesis of Methionine-Derived Glucosinolates in *Arabidopsis thaliana*. The Plant Cell, 21(6), 1813–1829. doi: 10.1105/tpc.109.066399

Gil, S. S., Cappellari, L. del R., Giordano, W., & Banchio, E. 2023. Antifungal Activity and Alleviation of Salt Stress by Volatile Organic Compounds of Native Pseudomonas Obtained from Mentha piperita. Plants, 12(7), 1488. doi: 10.3390/plants12071488

Goh, T., Kasahara, H., Mimura, T., Kamiya, Y., & Fukaki, H. 2012. Multiple AUX/IAA–ARF modules regulate lateral root formation: the role of *Arabidopsis* SHY2/IAA3-mediated auxin signalling. Philosophical Transactions of the Royal Society B: Biological Sciences, 367(1595), 1461–1468. doi: 10.1098/rstb.2011.0232

González Ortega-Villaizán, A., King, E., Patel, M. K., et al. 2024. The endophytic fungus *Serendipita indica* affects auxin distribution in *Arabidopsis thaliana* roots through alteration of auxin transport and conjugation to promote plant growth. Plant, Cell & Environment, 47(10), 3899–3919. doi: 10.1111/pce.14989

Groenhagen, U., Baumgartner, R., Bailly, A., Gardiner, A., Eberl, L., Schulz, S., & Weisskopf, L. 2013. Production of Bioactive Volatiles by Different Burkholderia ambifaria Strains. Journal of Chemical Ecology, 39(7), 892–906. doi: 10.1007/s10886-013-0315-y

Guo, Y., Jud, W., Weikl, F., Ghirardo, A., Junker, R. R., Polle, A., Benz, J. P., Pritsch, K., Schnitzler, J.-P., & Rosenkranz, M. 2021. Volatile organic compound patterns predict fungal trophic mode and lifestyle. Communications Biology, 4(1), 673. doi: 10.1038/s42003-021-02198-8

Gupta, M., Wong, M., Jawed, K., et al. 2022. Isobutanol production by combined in vivo and in vitro metabolic engineering. Metabolic Engineering Communications, 15, e00210. doi: 10.1016/j.mec.2022.e00210

Hao, F., Zhao, S., Dong, H., Zhang, H., Sun, L., & Miao, C. 2010. Nia1 and Nia2 are Involved in Exogenous Salicylic Acid-induced Nitric Oxide Generation and Stomatal Closure in *Arabidopsis*. Journal of Integrative Plant Biology, 52(3), 298–307. doi: 10.1111/j.1744-7909.2010.00920.x

Higuchi, M., Pischke, M. S., Pekka Mähönen, A., et al. 2004. *In planta* functions of the Arabidopsis cytokinin receptor family. In PLANT BIOLOGY (Vol. 101, Issue 23). doi: 10.1073/pnas.0402887101

Hpoo, M. K., Mishyna, M., Prokhorov, V., Arie, T., Takano, A., Oikawa, Y., & Fujii, Y. 2020. Potential of Octanol and Octanal from Heracleum sosnowskyi Fruits for the Control of Fusarium oxysporum f. sp. lycopersici. Sustainability, 12(22), 9334. doi: 10.3390/su12229334

Hong, J. P., Takeshi, Y., Kondou, Y., Schachtman, D. P., Matsui, M., & Shin, R. 2013. Identification and Characterization of Transcription Factors Regulating Arabidopsis HAK5. Plant and Cell Physiology, 54(9), 1478–1490. doi: 10.1093/PCP/PCT094

Hung, R., Lee, S., & Bennett, J. W. 2015. Fungal volatile organic compounds and their role in ecosystems. Applied Microbiology and Biotechnology, 99(8), 3395–3405. doi: 10.1007/s00253-015-6494-4

Inamdar, A. A., Morath, S., & Bennett, J. W. 2020. Fungal Volatile Organic Compounds: More Than Just a Funky Smell? Annual Review of Microbiology, 74(1), 101–116. doi: 10.1146/annurev-micro-012420-080428

Inoue, T., Higuchi, M., Hashimoto, Y., Seki, M., Kobayashi, M., Kato, T., Tabata, S., Shinozaki, K., & Kakimoto, T. 2001. Identification of CRE1 as a cytokinin receptor from Arabidopsis. Nature, 409(6823), 1060–1063. doi: 10.1038/35059117

Jing, H., & Strader, L. 2019. Interplay of Auxin and Cytokinin in Lateral Root Development. International Journal of Molecular Sciences, 20(3), 486. doi: 10.3390/ijms20030486

Jung, S. C., Martinez-Medina, A., Lopez-Raez, J. A., & Pozo, M. J. 2012. Mycorrhiza-Induced Resistance and Priming of Plant Defenses. Journal of Chemical Ecology, 38(6), 651–664. doi: 10.1007/s10886-012-0134-6

Jung, J. Y., Shin, R., & Schachtman, D. P. 2009. Ethylene Mediates Response and Tolerance to Potassium Deprivation in Arabidopsis. The Plant Cell, 21(2), 607–621. doi: 10.1105/TPC.108.063099

Kazan, K. 2013. Auxin and the integration of environmental signals into plant root development. Annals of Botany, 112(9), 1655–1665. doi: 10.1093/aob/mct229

Kim, D., Langmead, B., & Salzberg, S. L. 2015. HISAT: a fast spliced aligner with low memory requirements. Nature Methods 2015 12:4, 12(4), 357–360. doi: 10.1038/nmeth.3317

Kolde R. 2018. pheatmap: Pretty Heatmaps. R package version 1.0.12, https://github.com/raivokolde/pheatmap. (n.d.).

Kroymann, J., Donnerhacke, S., Schnabelrauch, D., & Mitchell-Olds, T. 2003. Evolutionary dynamics of an Arabidopsis insect resistance quantitative trait locus. Proceedings of the National Academy of Sciences of the United States of America, 100(24), 14587–14592. doi: 10.1073/pnas.1734046100

Kurepa, J., Shull, T. E., & Smalle, J. A. 2019. Antagonistic activity of auxin and cytokinin in shoot and root organs. Plant Direct, 3(2). doi: 10.1002/pld3.121

Kurepa, J., & Smalle, J. A. 2022. Auxin/Cytokinin Antagonistic Control of the Shoot/Root Growth Ratio and Its Relevance for Adaptation to Drought and Nutrient Deficiency Stresses. International Journal of Molecular Sciences, 23(4), 1933. doi: 10.3390/ijms23041933

Laller, R., Khosla, P. K., Negi, N., Avinash, H., Kusum, Thakur N., Kashyap, S., Shukla, S. K., & Hussain, I. 2023. Bacterial volatiles as PGPRs: Inducing plant defense mechanisms during stress periods. South African Journal of Botany, 159, 131–139. doi: 10.1016/j.sajb.2023.05.041

Ledford, W. C., Silvestri, A., Fiorilli, V., Roth, R., Rubio-Somoza, I., & Lanfranco, L. 2024. A journey into the world of small RNAs in the arbuscular mycorrhizal symbiosis. New Phytologist, 242(4), 1534–1544. doi: 10.1111/NPH.19394

Le Hir, R., Spinner, L., Klemens, P. A. W., et al. 2015. Disruption of the Sugar Transporters *AtSWEET11* and *AtSWEET12* Affects Vascular Development and Freezing Tolerance in Arabidopsis. Molecular Plant, 8(11), 1687–1690. doi: 10.1016/j.molp.2015.08.007

Lee, S., Yap, M., Behringer, G., Hung, R., & Bennett, J. W. 2016. Volatile organic compounds emitted by Trichoderma species mediate plant growth. Fungal Biology and Biotechnology, 3(1), 7. doi: 10.1186/s40694-016-0025-7

Li, J., Wu, W. H., & Wang, Y. 2017. Potassium channel AKT1 is involved in the auxin-mediated root growth inhibition in Arabidopsis response to low K+ stress. Journal of Integrative Plant Biology, 59(12), 895–909. doi: 10.1111/JIPB.12575

Li, H., Handsaker, B., Wysoker, A., Fennell, T., Ruan, J., Homer, N., Marth, G., Abecasis, G., & Durbin, R. 2009. The Sequence Alignment/Map format and SAMtools. Bioinformatics, 25(16), 2078–2079. doi: 10.1093/BIOINFORMATICS/BTP352

Li, Q., Zhu, X., Xie, Y., & Liang, J. 2021. Antifungal properties and mechanisms of three volatile aldehydes (octanal, nonanal and decanal) on *Aspergillus flavus*. Grain & Oil Science and Technology, 4(3), 131–140. doi: 10.1016/J.GAOST.2021.07.002

Liao, Y., Smyth, G. K., & Shi, W. 2014. featureCounts: an efficient general purpose program for assigning sequence reads to genomic features. Bioinformatics, 30(7), 923–930. doi: 10.1093/BIOINFORMATICS/BTT656

Lichtenthaler, H. K. 1987. Chlorophylls end carotenoids: Pigments of photosynthetic bio membranes. Methods in Enzimilogy, 148: 350–382. doi: 10.1016/0076-6879(87)48036-1

Lincoln, C., Britton, J. H., & Estelle, M. 1990. Growth and development of the *axr1* mutants of Arabidopsis. The Plant Cell, 2(11), 1071–1080. doi: 10.1105/tpc.2.11.1071

Love, M. I., Huber, W., & Anders, S. 2014. Moderated estimation of fold change and dispersion for RNA-seq data with DESeq2. Genome Biology 2014 15:12, 15(12), 550. doi: 10.1186/S13059-014-0550-8

Lozano-Juste, J., & León, J. 2010. Nitric oxide modulates sensitivity to ABA. Plant Signaling and Behavior, 5(3), 314–316. doi:10.4161/psb.5.3.11235

Luschnig, C., Gaxiola, R. A., Grisafi, P., & Fink, G. R. 1998. EIR1, a root-specific protein involved in auxin transport, is required for gravitropism in *Arabidopsis thaliana*. Genes & Development, 12(14), 2175–2187. doi: 10.1101/gad.12.14.2175

Mallick, A., Dey, S., Datta, S., Barman, M., Samui, S., & Dutta, G. 2022. Auxin and cytokinin signaling in plant stress response. In Auxins, Cytokinins and Gibberellins Signaling in Plants (pp. 213–234). Springer International Publishing. doi: 10.1007/978-3-031-05427-3_9

Marmeisse, R., Guidot, A., Gay, G., Lambilliotte, R., Sentenac, H., Combier, J.-P., Melayah, D., Fraissinet-Tachet, L., & Debaud, J. C. 2004. *Hebeloma cylindrosporum*–a model species to study ectomycorrhizal symbiosis from gene to ecosystem. New Phytologist, 163(3), 481–498. doi: 10.1111/j.1469-8137.2004.01148.x

Martínez-Santos, E., Cruz-Cruz, C. A., Spinoso-Castillo, J. L., & Bello-Bello, J. J. 2021. *In vitro* response of vanilla (*Vanilla planifolia Jacks. ex Andrews*) to PEG-induced osmotic stress. Scientific Reports, 11(1), 22611. doi: 10.1038/s41598-021-02207-0

Miñambres, E., Chaparro-Arias, M., Señorans, J., Valera-León, S., Soria-Solabarrieta, A., & Calvo-Polanco, M. 2026. Mycorrhizal fungi volatiles: determining the fate of plants against stress? Frontiers in Plant Science, 17, 1756587. doi: 10.3389/FPLS.2026.1756587

Moisan, K., Raaijmakers, J. M., Dicke, M., Lucas-Barbosa, D., & Cordovez, V. 2021. Volatiles from soil-borne fungi affect directional growth of roots. Plant, Cell & Environment, 44(1), 339–345. doi: 10.1111/pce.13890

Mravec, J., Skůpa, P., Bailly, A., et al. 2009. Subcellular homeostasis of phytohormone auxin is mediated by the ER-localized PIN5 transporter. Nature, 459(7250), 1136–1140. doi: 10.1038/nature08066

Murashige, T. 1962. A revised medium for rapid growth and bio assays with tobacco tissue cultures. Physiologia plantarum, 15(3). doi: 10.1111/j.1399-3054.1962.tb08052.x

Naser, V., & Shani, E. 2016. Auxin response under osmotic stress. Plant Molecular Biology 2016 91:6, 91(6), 661–672. doi: 10.1007/S11103-016-0476-5

Nenadić, M., & Vermeer, J. E. M. 2021. Dynamic cytokinin signalling landscapes during lateral root formation in Arabidopsis. Quantitative Plant Biology, 2, e13. doi: 10.1017/QPB.2021.13

O’Callaghan, M., Ballard, R. A., & Wright, D. 2022. Soil microbial inoculants for sustainable agriculture: Limitations and opportunities. Soil Use and Management, 38(3), 1340–1369. doi: 10.1111/sum.12811

Olas, J. J., & Wahl, V. 2019. Tissue-specific *NIA1* and *NIA2* expression in *Arabidopsis thaliana*. Plant Signaling & Behavior, 14(11), 1656035. doi: 10.1080/15592324.2019.1656035

Opitz, M. W., Daneshkhah, R., Lorenz, C., Ludwig, R., Steinkellner, S., & Wieczorek, K. 2021. *Serendipita indica* changes host sugar and defense status in *Arabidopsis thaliana*: cooperation or exploitation? Planta, 253(3), 74. doi: 10.1007/s00425-021-03587-3

Pachlewski, R., & Pachlewska, J. 1974. Studies on symbiotic properties of mycorrhizal fungi of pine (*Pinus silvestris L*.) with the aid of the method of mycorrhizal synthesis in pure cultures on agar. Forest Research Institute.

Pagès H., Carlson M., Falcon S., Li N. 2024. AnnotationDbi: Manipulation of SQLite-based annotations in Bioconductor. R package version 1.68.0, https://bioconductor.org/packages/AnnotationDbi. (n.d.).

Pérez-Flores, P., Valencia-Cantero, E., Altamirano-Hernández, J., Pelagio-Flores, R., López-Bucio, J., García-Juárez, P., & Macías-Rodríguez, L. 2017. *Bacillus methylotrophicus* M4-96 isolated from maize (*Zea mays*) rhizoplane increases growth and auxin content in *Arabidopsis thaliana* via emission of volatiles. Protoplasma, 254(6), 2201–2213. doi: 10.1007/s00709-017-1109-9

Perilli, S., Moubayidin, L., & Sabatini, S. 2010. The molecular basis of cytokinin function. Current Opinion in Plant Biology, 13(1), 21–26. doi: 10.1016/j.pbi.2009.09.018

Petersen, B., Chen, S., Hansen, C., Olsen, C., & Halkier, B. 2002. Composition and content of glucosinolates in developing *Arabidopsis thaliana*. Planta, 214(4), 562–571. doi: 10.1007/s004250100659

Pickett, F. B., Wilson, A. K., & Estelle, M. 1990. The *aux1* Mutation of *Arabidopsis* Confers Both Auxin and Ethylene Resistance. Plant Physiology, 94(3), 1462–1466. doi: 10.1104/pp.94.3.1462

Plett, J. M., Wojtalewicz, D., Plett, K. L., Collin, S., Kohler, A., Jacob, C., & Martin, F. 2024. Sesquiterpenes of the ectomycorrhizal fungus *Pisolithus microcarpus* alter root growth and promote host colonization. Mycorrhiza 2024 34:1, 34(1), 69–84. doi: 10.1007/S00572-024-01137-9

Pons, S., Fournier, S., Chervin, C., Bécard, G., Rochange, S., Dit Frey, N. F., & Pagès, V. P. 2020. Phytohormone production by the arbuscular mycorrhizal fungus *Rhizophagus irregularis*. PLOS ONE, 15(10), e0240886. doi: 10.1371/JOURNAL.PONE.0240886

Poschet, G., Hannich, B., & Büttner, M. 2010. Identification and Characterization of *AtSTP14*, a Novel Galactose Transporter from Arabidopsis. Plant and Cell Physiology, 51(9), 1571–1580. doi: 10.1093/pcp/pcq100

Quint, M., & Gray, W. M. 2006. Auxin signaling. Current Opinion in Plant Biology, 9(5), 448–453. doi: 10.1016/j.pbi.2006.07.006

Quint, M., Ito, H., Zhang, W., & Gray, W. M. 2005. Characterization of a novel temperature-sensitive allele of the *CUL1/AXR6* subunit of SCF ubiquitin-ligases. The Plant Journal, 43(3), 371–383. doi: 10.1111/j.1365-313X.2005.02449.x

Ravelo-Ortega, G., López-Bucio, J. S., Ruiz-Herrera, L. F., Pelagio-Flores, R., Ayala-Rodríguez, J. Á., de la Cruz, H. R., Guevara-García, Á. A., & López-Bucio, J. 2021. The growth of Arabidopsis primary root is repressed by several and diverse amino acids through auxin-dependent and independent mechanisms and MPK6 kinase activity. Plant Science, 302, 110717. doi: 10.1016/j.plantsci.2020.110717

Rawat, S. S., & Laxmi, A. 2024. Light at the end of the tunnel: integrating signaling pathways in the coordination of lateral root development. Biochemical Society Transactions, 52(4), 1895–1908. doi: 10.1042/BST20240049

R Core Team. 2024. R: A language and environment for statistical computing [Software]. R Foundation for Statistical Computing. https://www.R-project.org/. (n.d.).

Rubio, F., Nieves-Cordones, M., Alemán, F., & Martínez, V. 2008. Relative contribution of *AtHAK5* and *AtAKT1* to K+ uptake in the high-affinity range of concentrations. Physiologia Plantarum, 134(4), 598–608. doi: 10.1111/j.1399-3054.2008.01168.x

Ruytinx, J., Miyauchi, S., Hartmann-Wittulsky, S., et al. 2021. A Transcriptomic Atlas of the Ectomycorrhizal Fungus *Laccaria bicolor*. Microorganisms, 9(12), 2612. doi: 10.3390/microorganisms9122612

Sánchez-López, Á. M., Baslam, M., de Diego, N., et al. 2016. Volatile compounds emitted by diverse phytopathogenic microorganisms promote plant growth and flowering through cytokinin action. Plant, Cell & Environment, 39(12), 2592–2608. doi: 10.1111/pce.12759

Santos Teixeira, J. A., & ten Tusscher, K. H. 2019. The Systems Biology of Lateral Root Formation: Connecting the Dots. Molecular Plant, 12(6), 784–803. doi: 10.1016/J.MOLP.2019.03.015

Schneider, C. A., Rasband, W. S., & Eliceiri, K. W. 2012. NIH Image to ImageJ: 25 years of image analysis. Nature Methods, 9(7), 671–675. doi: 10.1038/nmeth.2089

Schnürer, J., Olsson, J., & Börjesson, T. 1999. Fungal Volatiles as Indicators of Food and Feeds Spoilage. Fungal Genetics and Biology, 27(2–3), 209–217. doi: 10.1006/fgbi.1999.1139

Schuster, J., Knill, T., Reichelt, M., Gershenzon, J., & Binder, S. 2006. BRANCHED-CHAIN AMINOTRANSFERASE4 Is Part of the Chain Elongation Pathway in the Biosynthesis of Methionine-Derived Glucosinolates in Arabidopsis. The Plant Cell, 18(10), 2664–2679. doi: 10.1105/tpc.105.039339

Skoog, F., & Miller, C. O. 1957. Chemical regulation of growth and organ formation in plant tissues cultured *in vitro*. Symp Soc Exp Biol, 11, 118–130.

Smith, S. E., & Read, D. J. (2010). Mycorrhizal symbiosis. Academic press. doi: 10.1016/B978-0-12-370526-6.X5001-6

Tewari, R. K., Horemans, N., & Watanabe, M. 2021. Evidence for a role of nitric oxide in iron homeostasis in plants. Journal of Experimental Botany, 72(4), 990–1006. doi: 10.1093/jxb/eraa484

Tian, Q., & Reed, J. W. 1999. Control of auxin-regulated root development by the *Arabidopsis thaliana SHY2/IAA3* gene. Development, 126(4), 711–721. doi: 10.1242/dev.126.4.711

Tyagi, J., Chaudhary, P., Mishra, A., Khatwani, M., Dey, S., & Varma, A. 2022. Role of Endophytes in Abiotic Stress Tolerance: With Special Emphasis on *Serendipita indica*. International Journal of Environmental Research, 16(4), 62. doi: 10.1007/s41742-022-00439-0

Uddin, M. N., Rahman, U. ur, khan, W., Uddin, N., & Muhammad, M. 2018. Effect of *Trichoderma harzianum* on tomato plant growth and its antagonistic activity against *Phythium ultimum* and *Phytopthora capsici*. Egyptian Journal of Biological Pest Control, 28(1), 32. doi: 10.1186/s41938-018-0032-5

Vahabi, K., Camehl, I., Sherameti, I., & Oelmüller, R. 2013. Growth of Arabidopsis seedlings on high fungal doses of *Piriformospora indica* has little effect on plant performance, stress, and defense gene expression in spite of elevated jasmonic acid and jasmonic acid-isoleucine levels in the roots. Plant Signaling & Behavior, 8(11). doi: 10.4161/psb.26301

Vahabi, K., Sherameti, I., Bakshi, M., Mrozinska, A., Ludwig, A., Reichelt, M., & Oelmüller, R. 2015. The interaction of Arabidopsis with *Piriformospora indica* shifts from initial transient stress induced by fungus-released chemical mediators to a mutualistic interaction after physical contact of the two symbionts. BMC Plant Biology 2015 15:1, 15(1), 58-. doi: 10.1186/s12870-015-0419-3

Valmas, M. I., Sexauer, M., Markmann, K., & Tsikou, D. 2023. Plants Recruit Peptides and Micro RNAs to Regulate Nutrient Acquisition from Soil and Symbiosis. Plants, 12(1), 187. doi: 10.3390/plants12010187

van der Does, D., Boutrot, F., Engelsdorf, T., et al. 2017. The Arabidopsis leucine-rich repeat receptor kinase MIK2/LRR-KISS connects cell wall integrity sensing, root growth and response to abiotic and biotic stresses. PLOS Genetics, 13(6), e1006832. doi: 10.1371/journal.pgen.1006832

Valenzuela-Aragon, B., Cardinale, M., Rolli, E., Rustioni, L., & Francioli, D. 2025. The role of arbuscular mycorrhizal fungi in abiotic stress management in viticulture under climatic shifts. Plant Stress, 16, 100863. doi: 10.1016/J.STRESS.2025.100863

Varma, A., Savita Verma, Sudha, Sahay, N., Bütehorn, B., & Franken, P. 1999. *Piriformospora indica*, a Cultivable Plant-Growth-Promoting Root Endophyte. Applied and Environmental Microbiology, 65(6), 2741–2744. doi: 10.1128/AEM.65.6.2741-2744.1999

Vazquez, M. M., Casalongué, C. A., & París, R. 2019. Nitrate reductase mediates nitric oxide-dependent gravitropic response in *Arabidopsis thaliana* roots. Plant Signaling & Behavior, 14(4), e1578631. doi: 10.1080/15592324.2019.1578631

Verma, S., Varma, A., Rexer, K.-H., Hassel, A., Kost, G., Sarbhoy, A., Bisen, P., Bütehorn, B., & Franken, P. 1998. *Piriformospora indica*, gen. et sp. nov., a new root-colonizing fungus. Mycologia, 90(5), 896–903. doi: 10.1080/00275514.1998.12026983

Venneman, J., Vandermeersch, L., Walgraeve, C., Audenaert, K., Ameye, M., Verwaeren, J., Steppe, K., van Langenhove, H., Haesaert, G., & Vereecke, D. 2020. Respiratory CO2 Combined With a Blend of Volatiles Emitted by Endophytic *Serendipita* Strains Strongly Stimulate Growth of Arabidopsis Implicating Auxin and Cytokinin Signaling. Frontiers in Plant Science, 11, 544435. doi: 10.3389/FPLS.2020.544435

Vert, G., Grotz, N., Dédaldéchamp, F., Gaymard, F., Guerinot, M. L., Briat, J. F., & Curie, C. 2021. CORRECTION: IRT1, an arabidopsis transporter essential for iron uptake from the soil and for plant growth. The Plant Cell, 33(2), 439–440. doi: 10.1093/plcell/koaa033

Wang, J., Yu, X., Ding, Z. J., et al. 2021. Structural basis of ALMT1-mediated aluminum resistance in Arabidopsis. Cell Research 2021 32:1, 32(1), 89–98. doi: 10.1038/s41422-021-00587-6

Werner, T., & Schmülling, T. 2009. Cytokinin action in plant development. Current Opinion in Plant Biology, 12(5), 527–538. doi: 10.1016/j.pbi.2009.07.002

Wickham, H. 2016. ggplot2. Springer International Publishing. doi: 10.1007/978-3-319-24277-4

Wilkinson, J. Q., & Crawford, N. M. 1993. Identification and characterization of a chlorate-resistant mutant of *Arabidopsis thaliana* with mutations in both nitrate reductase structural genes *NIA1* and *NIA2*. MGG Molecular & General Genetics, 239(1–2), 289–297. doi: 10.1007/BF00281630

Wu, Q.-S., Zou, Y.-N., & Huang, Y.-M. 2013. The arbuscular mycorrhizal fungus *Diversispora spurca* ameliorates effects of waterlogging on growth, root system architecture and antioxidant enzyme activities of citrus seedlings. Fungal Ecology, 6(1), 37–43. doi: 10.1016/j.funeco.2012.09.002

Yasmin, H., Rashid, U., Hassan, M. N., Nosheen, A., Naz, R., Ilyas, N., Sajjad, M., Azmat, A., & Alyemeni, M. N. 2021. Volatile organic compounds produced by *Pseudomonas pseudoalcaligenes* alleviated drought stress by modulating defense system in maize (*Zea mays* L.). Physiologia Plantarum, 172(2), 896–911. doi: 10.1111/ppl.13304

Yokoyama, A., Yamashino, T., Amano, Y.-I., Tajima, Y., Imamura, A., Sakakibara, H., & Mizuno, T. 2006. Type-B ARR Transcription Factors, ARR10 and ARR12, are Implicated in Cytokinin-Mediated Regulation of Protoxylem Differentiation in Roots of *Arabidopsis thaliana*. Plant and Cell Physiology, 48(1), 84–96. doi: 10.1093/pcp/pcl040

Yu, G., Wang, L.-G., Han, Y., & He, Q.-Y. 2012. clusterProfiler: an R Package for Comparing Biological Themes Among Gene Clusters. OMICS: A Journal of Integrative Biology, 16(5), 284–287. doi: 10.1089/omi.2011.0118

Zeng, W., Xiang, D., Li, X., et al. 2025. Effects of combined inoculation of arbuscular mycorrhizal fungi and plant growth-promoting rhizosphere bacteria on seedling growth and rhizosphere microecology. Frontiers in Microbiology, 15. doi: 10.3389/fmicb.2024.1475485

Zhao, C., Zayed, O., Yu, Z., Jiang, W., Zhu, P., Hsu, C.-C., Zhang, L., Tao, W. A., Lozano-Durán, R., & Zhu, J.-K. 2018. Leucine-rich repeat extensin proteins regulate plant salt tolerance in *Arabidopsis*. Proceedings of the National Academy of Sciences, 115(51), 13123–13128. doi: 10.1073/pnas.1816991115

Zhao, S., Zhang, M. L., Ma, T. L., & Wang, Y. 2016. Phosphorylation of ARF2 Relieves Its Repression of Transcription of the K+ Transporter Gene HAK5 in Response to Low Potassium Stress. The Plant Cell, 28(12), 3005–3019. doi: 10.1105/TPC.16.00684

Zhu, X. F., Wang, B., Song, W. F., Zheng, S. J., & Shen, R. F. 2015. Putrescine Alleviates Iron Deficiency via NO-Dependent Reutilization of Root Cell-Wall Fe in Arabidopsis. Plant Physiology, 170(1), 558–567. doi: 10.1104/pp.15.01617

Zwiazek, J. J., Equiza, M. A., Karst, J., Senorans, J., Wartenbe, M., & Calvo-Polanco, M. 2019. Role of urban ectomycorrhizal fungi in improving the tolerance of lodgepole pine (*Pinus contorta*) seedlings to salt stress. Mycorrhiza 2019 29:4, 29(4), 303–312. doi: 10.1007/S00572-019-00893-3

