## Supplementary Tables for "Fungal volatiles affect lateral root formation and increase plant tolerance to osmotic stress"

**Supplementary Table 1. Oligonucleotides employed, including their nucleotide sequences and melting temperatures (Tm).** The *Elongation Factor 1* (*EF1*) and *Ubiquitin 5* (*UBQ5*) genes were used as internal housekeeping references.

| Gen | Name | Forward/Reverse | % GC | Tm (ºC) | 5’→ 3' |
|---|---|---|---|---|---|
| AT5G60390 | *EF1* | F | 47,8 | 63,0 | TGTAACAAGATGGATGCCACCAC |
| AT5G60390 | *EF1* | R | 54,5 | 64,0 | TCCCTCGAATCCAGAGATTGGC |
| AT3G62250 | *UBQ5* | F | 45,0 | 58,0 | CCAAGCCGAAGAAGATCAAG |
| AT3G62250 | *UBQ5* | R | 50,0 | 58,0 | ACTCCTTCCTCAAACGCTGA |
| AT4G13420 | *HAK5* | F | 60,0 | 58,3 | CTCTCGGCCAAAACACGGTG |
| AT4G13421 | *HAK5* | R | 60,0 | 58,0 | ATACGAGGATGATCGGGGCG |
| AT3G19710 | *BCAT4* | F | 55,0 | 57,7 | TAACCCTTGCGCTGCTGTTC |
| AT3G19710 | *BCAT4* | R | 60,0 | 58,0 | GAAGGCGGAGACCGTTTTGG |
| AT5G23660 | *SWEET12* | F | 55,0 | 57,7 | TTCGCATCCAAGAAAGCCCG |
| AT5G23660 | *SWEET12* | R | 55,0 | 57,4 | ATCTTCGCACGTGTGGTTCC |
| AT1G04240 | *SHY2/IAA3* | F | 50,0 | 57,7 | GAAAACTGAAACATCCCCTCCTCG |
| AT1G04240 | *SHY2/IAA3* | R | 50,0 | 58,5 | GATTCCTTGACCCTCATGCTCAGA |
| AT5G57090 | *EIR1/PIN2* | F | 65 | 60,2 | CCTCCTCACGACAACCTCGC |
| AT5G57090 | *EIR1/PIN2* | R | 60,0 | 59,1 | CGTCTTCCACGTCGCTACCT |
| AT2G38120 | *AUX1* | F | 60,0 | 58,3 | CTGTCTGGGACGCTTGGTTC |
| AT2G38120 | *AUX1* | R | 50,0 | 57,2 | GATGAGATAAGCAGTCCAGCTTCC |
| AT3G63110 | *IPT3* | F | 55,0 | 59,2 | CGCAACCGCTGTTGAAGCTT |
| AT3G63110 | *IPT3* | R | 65 | 60,2 | GCTAACGTGGCCTGGGAGAG |
| AT1G19770 | *PUP14* | F | 54,5 | 64 | GCTTATAGCCGGTGAGCAACAC |
| AT1G19770 | *PUP14* | R | 59,1 | 64 | ATCCCAACCCAATAGACCTGCC |
