## Supplementary Figures for "Fungal volatiles affect lateral root formation and increase plant tolerance to osmotic stress"

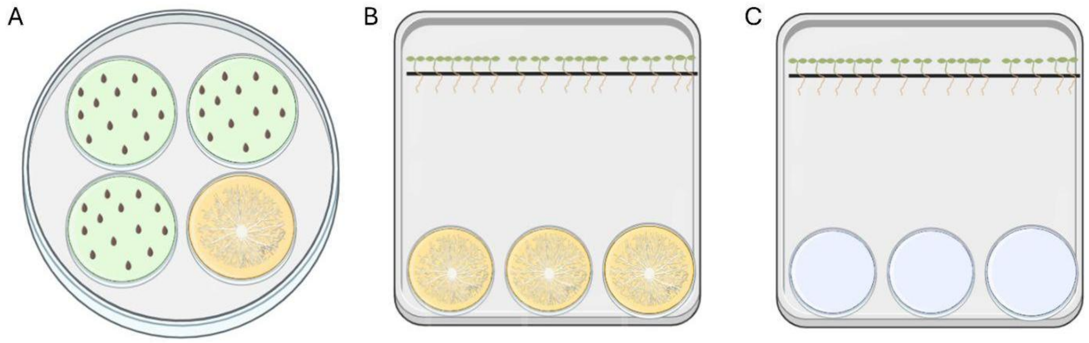


**Supplementary Figure 1. Experimental set-ups for the application of fungal VOCs.** A. Co-inoculation set-up of seedlings and fungi in the screening experiment with Col-0 and Arabidopsis mutants. B. Co-inoculation set-up of seedlings and fungi. C. Experimental set-up for the application of commercial VOCs. Created with BioRender.com.

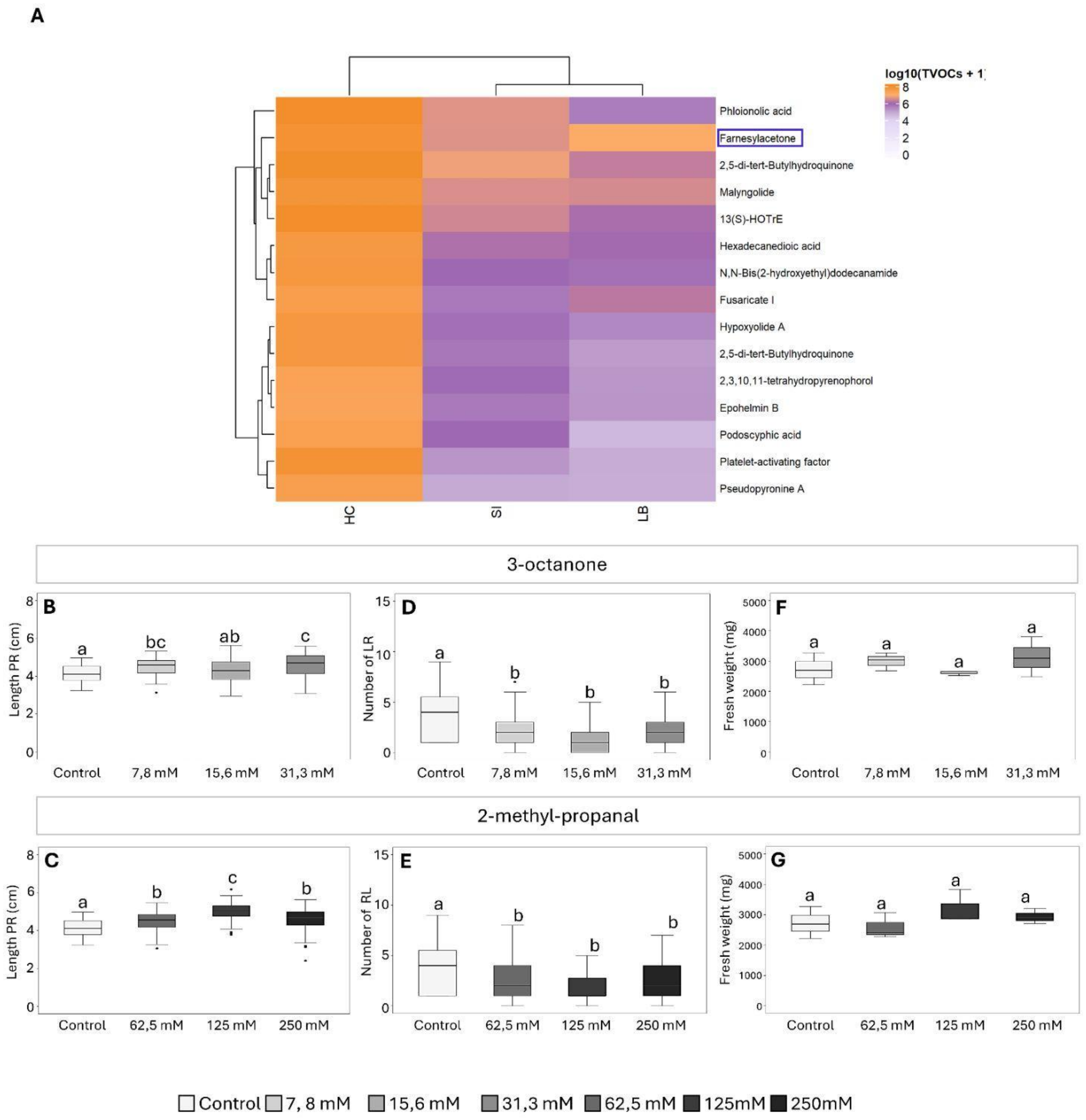


**Supplementary Figure 2. Profile of the compounds at the growing media after 6 days of exposure to fungal VOCs, and analysis of Col-0 10 days-old seedling in the presence of selected volatile compounds.** A. Heatmap of VOCs fixed in the seedlings growing media. Color intensity indicates the relative abundance of each compound in each condition, with rows representing individual volatiles and columns representing treatments. B-C. Primary root length (PR) (cm). D-E. Number of lateral roots (LR). F-G. Total fresh weight of seedlings (mg). Data are shown as boxplots of five biological replicates (n=5 plates) per genotype and condition; each plate contained 20 seeds per genotype. Different letters indicate statistical differences ($p<0.05$) among treatment means using one-way ANOVA and Tukey's test.

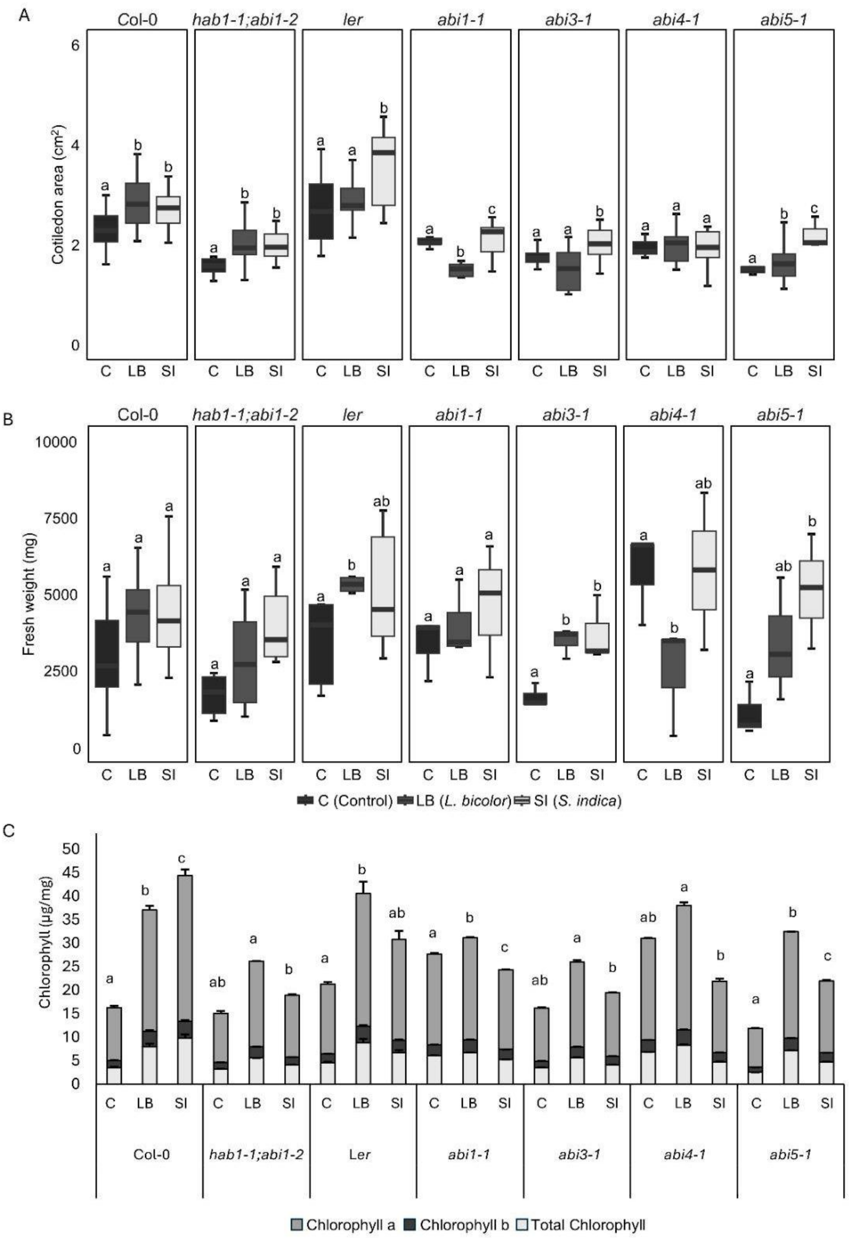


**Supplementary Figure 3. Seedling development of abscisic acid mutants (*hab1-1;abi1-2, abi1-1, abi3-1, abi4-1, abi5-1*) in the absence (C) or presence of VOCs emitted by *L. bicolor* (LB) and *S. indica* (SI).** A. Cotyledon area of 6-day-old seedlings ($cm^2$). B. Fresh weight of10 days-old seedling (mg). C. Total chlorophyll, chlorophyll a, and chlorophyll b (µg/mL) of 10 days-old seedlings. Three biological replicates (n=3 plates) per genotype and condition; each plate contained 60 seedlings. For chlorophyll measurements, five seedlings were pooled per biological replicate. Different letters indicate significant differences among treatment means according to one-way ANOVA followed by Tukey's test ($P < 0.05$).

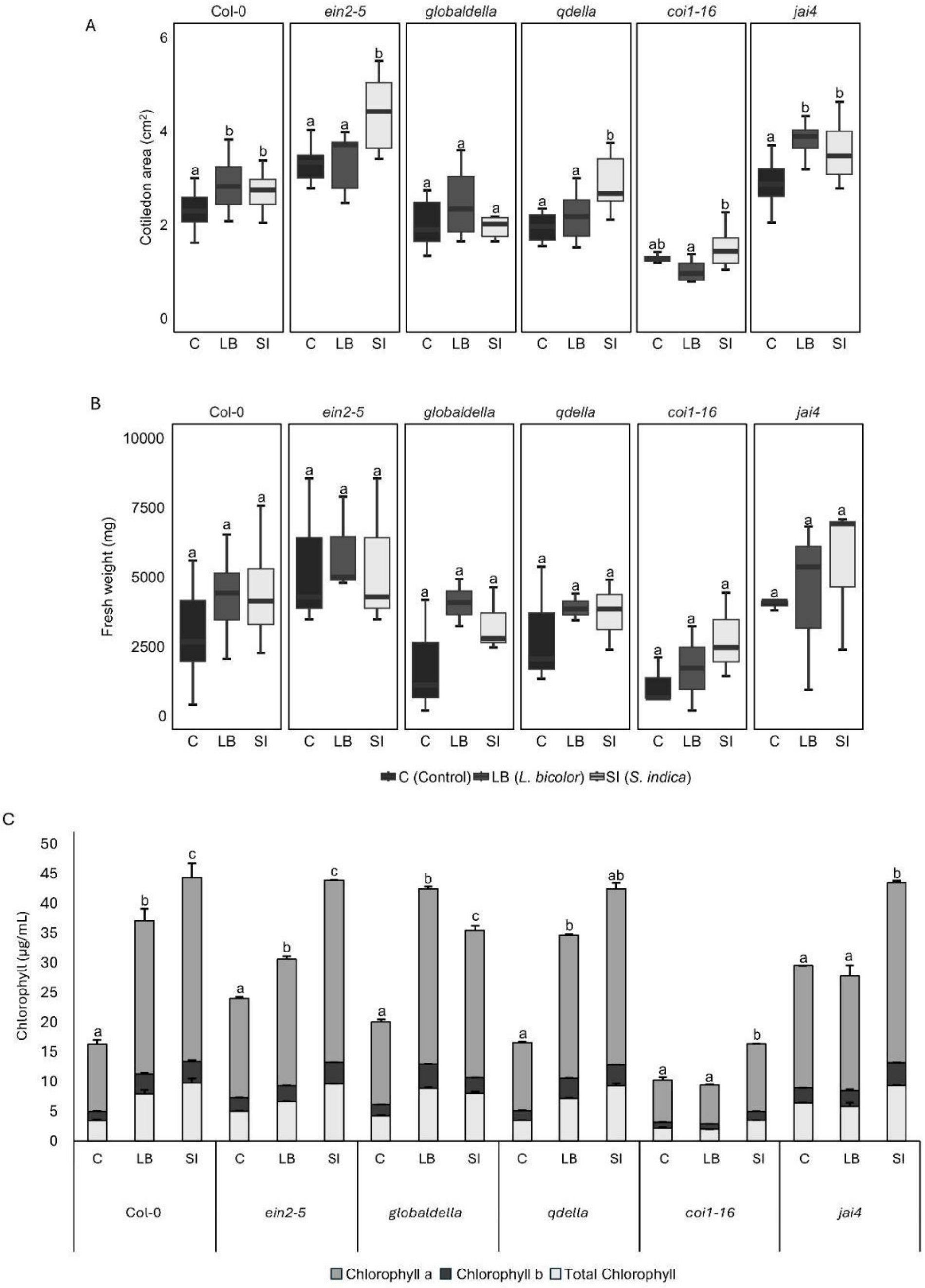


**Supplementary Figure 4. Development of the ethylene mutant (*ein2-5*), gibberellins mutants (*globaldella, qdella*) and jasmonic acid mutants (*coi1-16, jai4*) in the absence (C) or presence of VOCs emitted by *L. bicolor* (LB) and *S. indica* (SI).** A. Cotyledon area of 6-day-old seedlings ($cm^2$). B. Fresh weight of 10 days-old seedlings (mg). C. Total chlorophyll, chlorophyll a, and chlorophyll b (µg/mL) of 10 days-old seedlings. Three biological replicates (n=3 plates) per genotype and condition; each plate contained 60 seedlings. For chlorophyll measurements, five seedlings were pooled per biological replicate. Different letters indicate significant differences among treatment means according to one-way ANOVA followed by Tukey's test (P < 0.05).

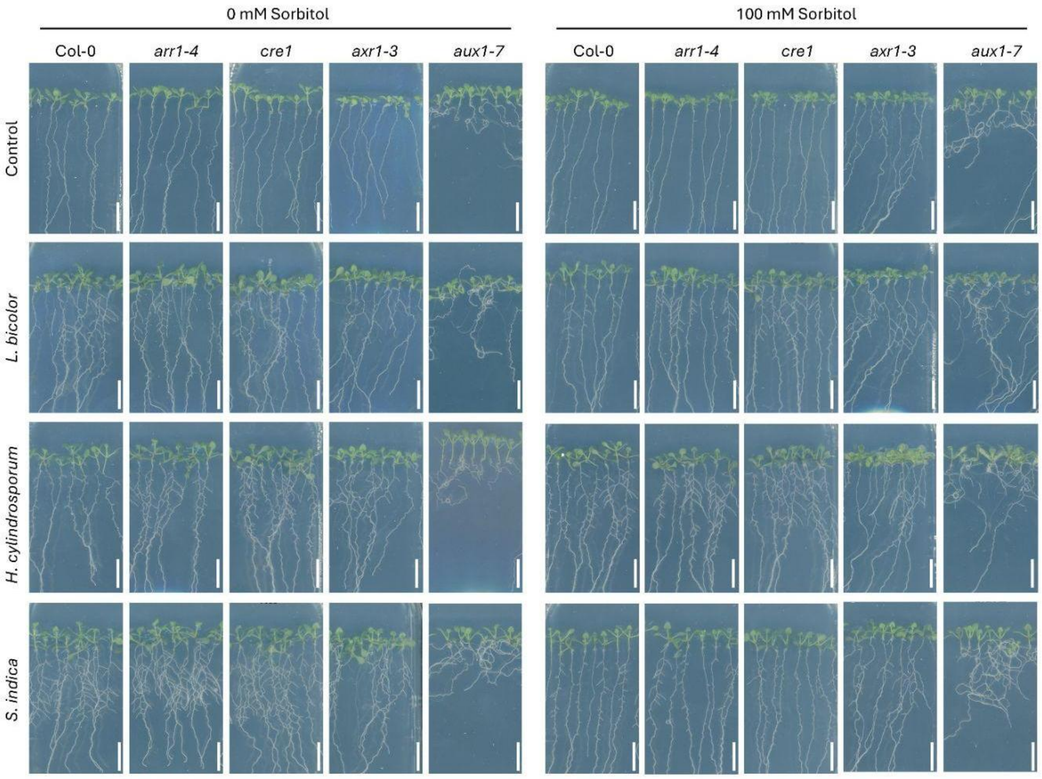


**Supplementary Figure 5. Phenotype of 10-day-old Arabidopsis seedlings in the absence and presence of VOCs from *L. bicolor*, *H. cylindrosporum*, and *S. indica.*** A. Control conditions (0 mM sorbitol). B. Osmotic stress conditions (100 mM sorbitol). Col-0 and the mutants *arr1-4*, *cre1*, *axr1-3*, and *aux1-7* are shown. Scale bar = 1 cm.

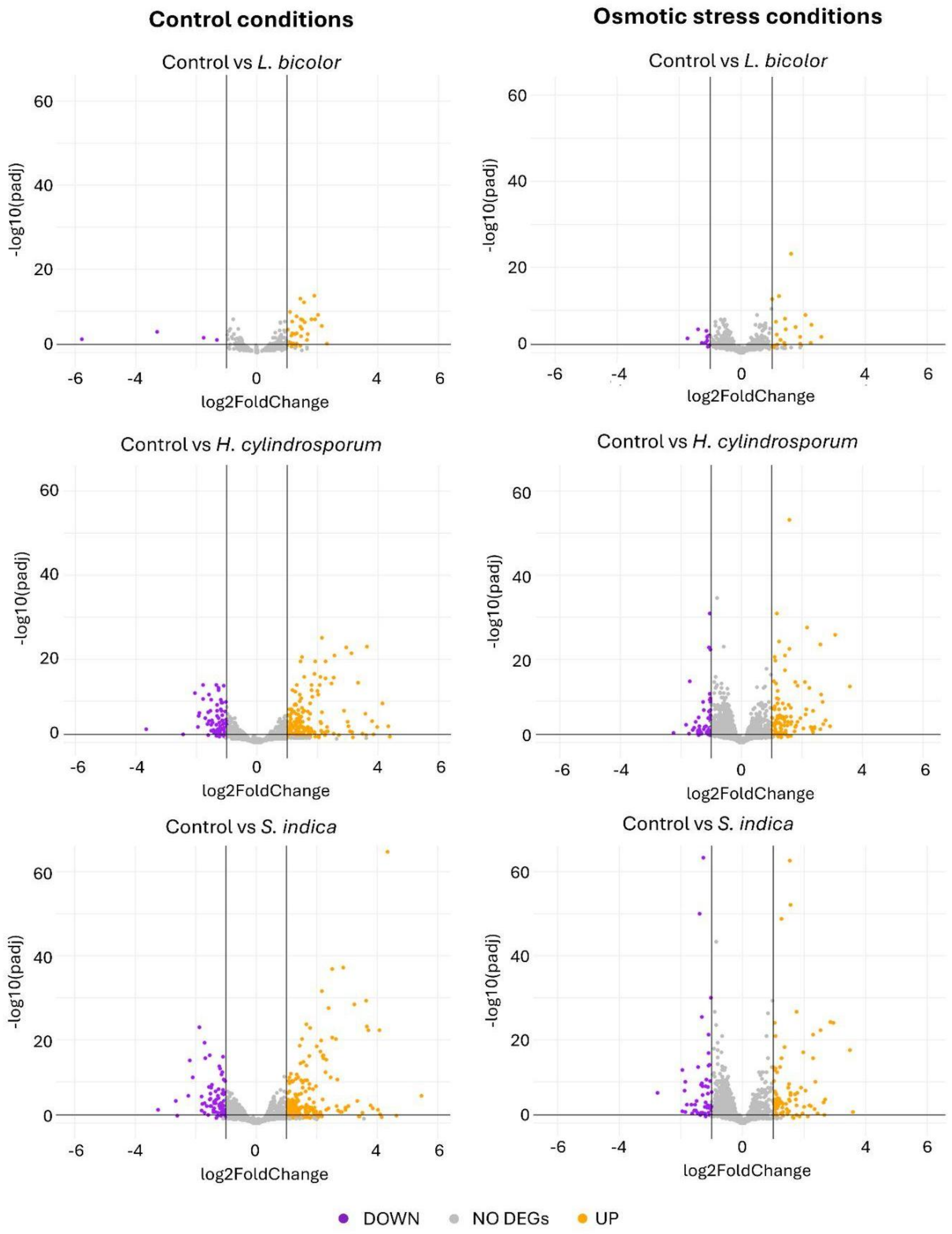


**Supplementary Figure 6. Volcano plots of differentially expressed genes (DEGs) in 10-day-old Col-0 seedlings exposed to VOCs emitted by *L. bicolor*, *H. cylindrosporum* or *S. indica*, under control conditions (0 mM sorbitol, left graphs) or osmotic stress conditions (100 mM sorbitol, right graphs).** Data are from three biological replicates (n=3) per condition. Purple dots indicate downregulated genes (DOWN), while the orange dots indicate upregulated genes (UP) (p-value $<0.05$, $\log_2$Foldchange$>|1.0|$).

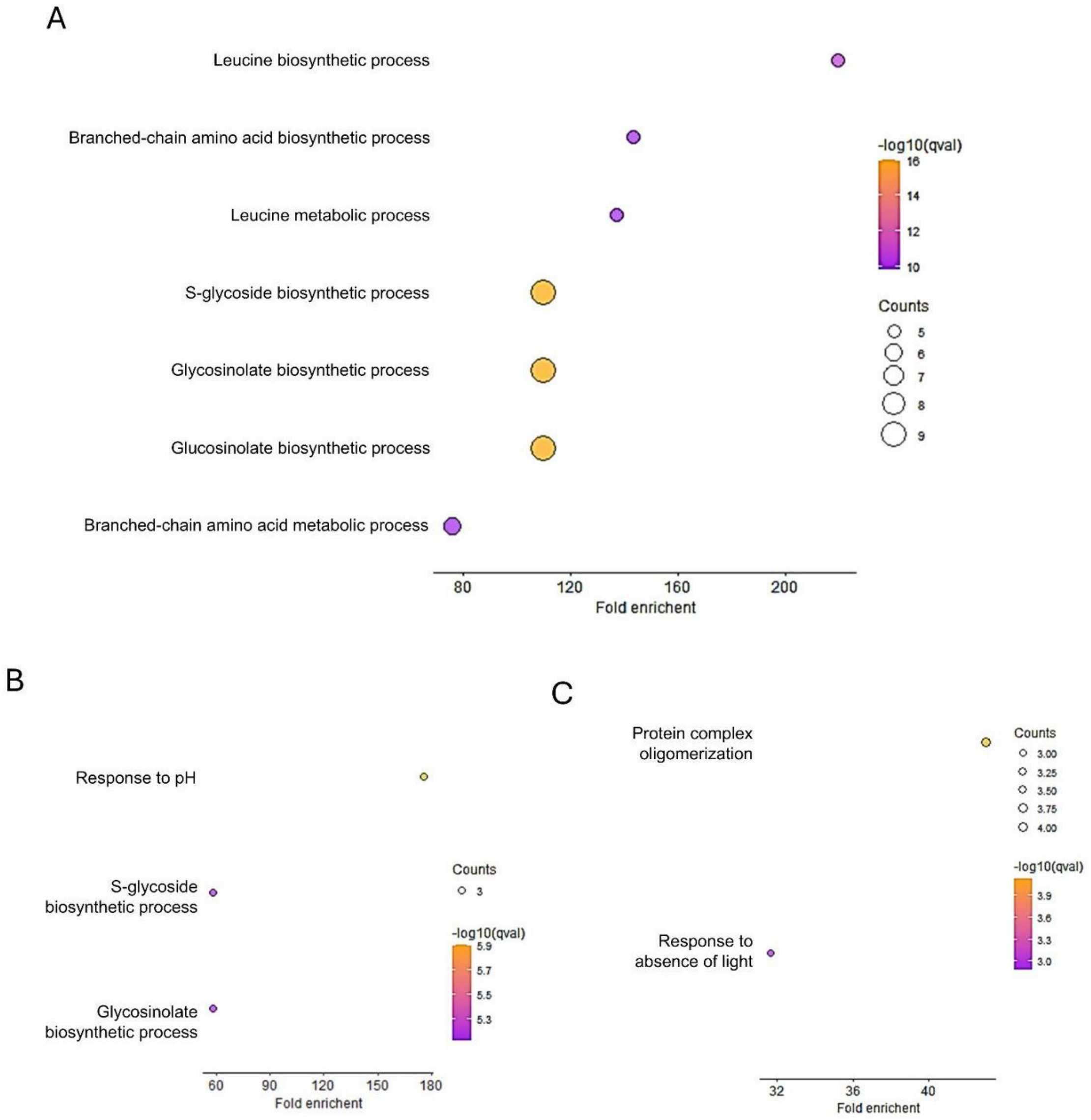


**Supplementary Figure 7. Ontological analysis of genes differentially expressed in roots of 10-day-old seedlings exposed to *L bicolor* VOCs.** A. Ontological analysis of genes induced under control conditions (0 mM sorbitol). B. Ontological analysis of genes induced under osmotic stress conditions (100 mM sorbitol). C. Ontological analysis of genes repressed under osmotic stress conditions (100 mM sorbitol).

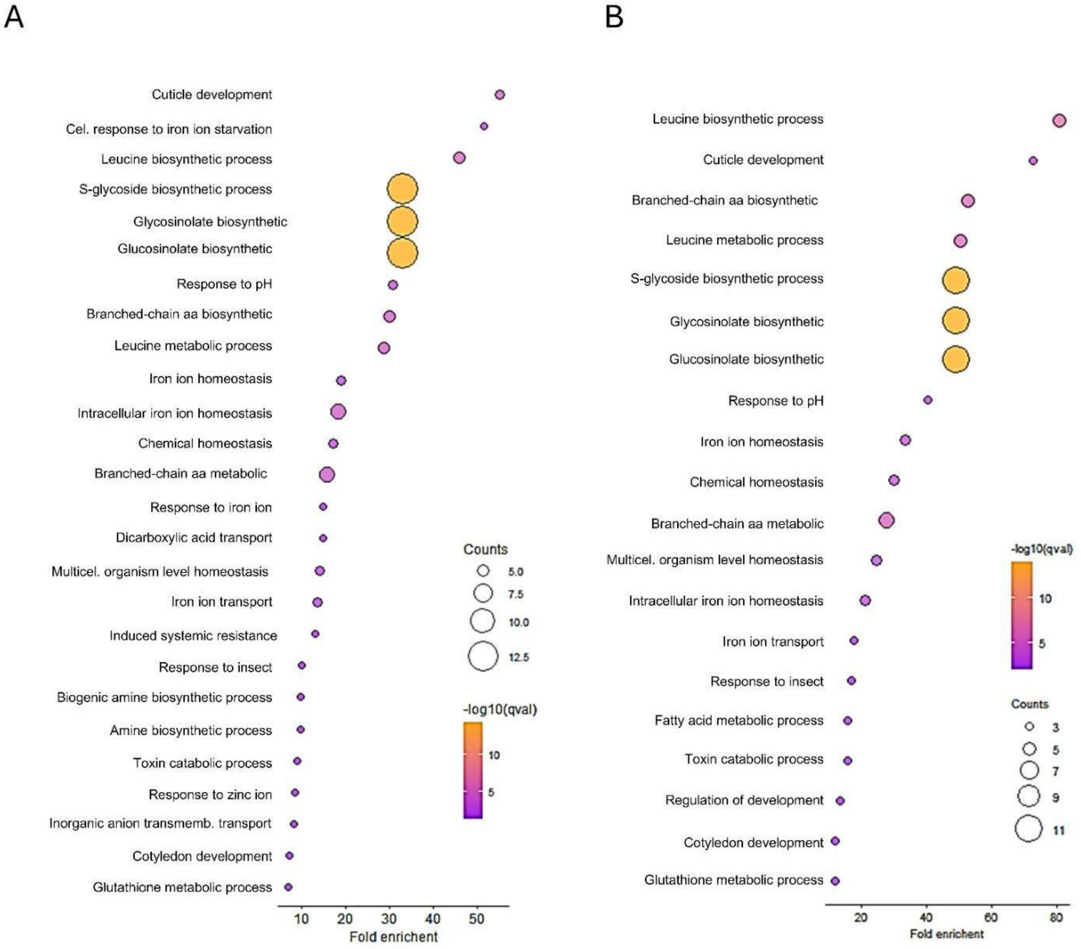


**Supplementary Figure 8. Ontological analysis of genes induced in roots of 10-day-old seedlings exposed to *H. cylindrosporum* VOCs.** A. Control conditions (0 mM Sorbitol). B. Osmotic stress conditions (100 mM Sorbitol).

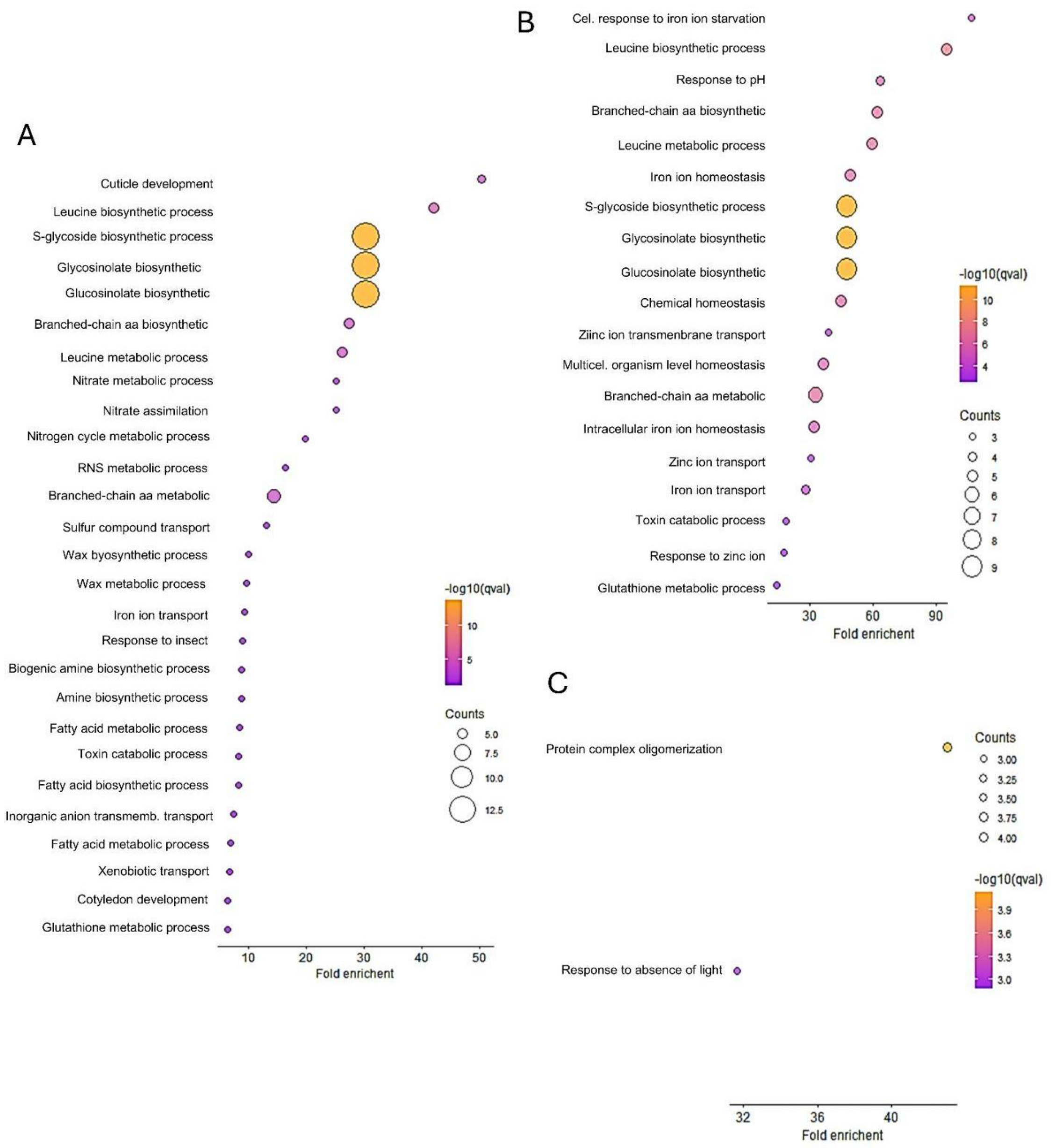


**Supplementary Figure 9. Ontological analysis of genes differentially expressed in roots of 10-day-old seedlings exposed to *S. indica* VOCs.** A. Ontological analysis of genes induced under control conditions (0 mM sorbitol). B. Ontological analysis of genes induced under osmotic stress conditions (100 mM Sorbitol). C. Ontological analysis of genes repressed under osmotic stress conditions (100 mM Sorbitol).

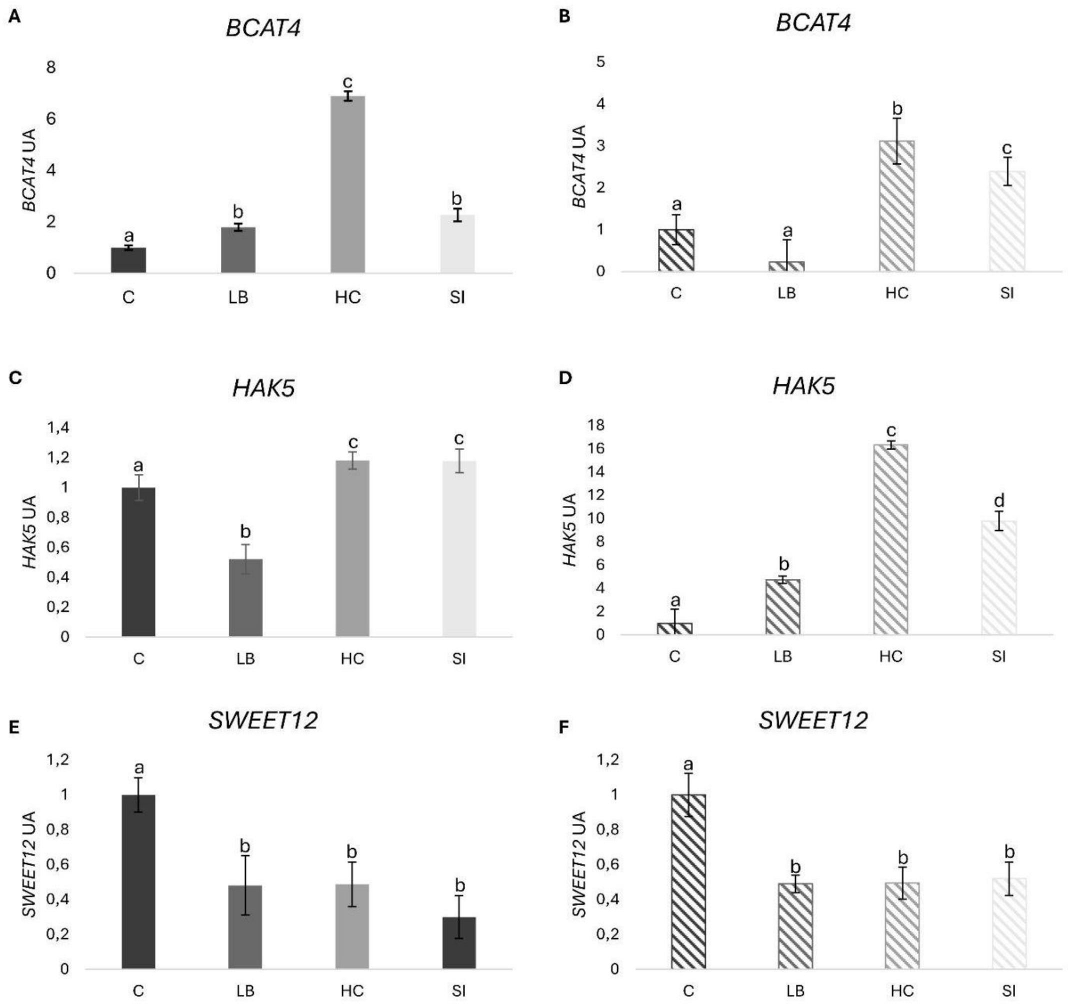


**Supplementary Figure 10. Gene expression of selected genes obtained from the transcriptomic profiling of 10-day-old Col-0 seedling roots exposed to the VOCs of *L. bicolor* (LB), *H. cylindrosporum* (HC), and *S. indica* (SI). The control condition (0 mM sorbitol) is depicted in solid color (left), whereas the osmotic stress condition (100 mM sorbitol) is shown in striped color (right).** A-B. *HAK5*. C-D. *BCAT*. E-F. *SWEET12*. Data are mean ± SE of three biological replicates (n = 3 independent plates), each analyzed in three technical replicates. Different letters indicate statistical differences ($p<0.05$) among treatment means using one-way ANOVA and Tukey's test.
